# KIDINS220 and InsP8 safeguard the stepwise regulation of phosphate exporter XPR1

**DOI:** 10.1101/2025.01.17.633679

**Authors:** Xiaojie Wang, Zhongjian Bai, Ciara Wallis, Huanchen Wang, Yaoyao Han, Ruitao Jin, Mingguang Lei, Tian Yang, Chunfang Gu, Henning Jessen, Stephen Shears, Yadong Sun, Ben Corry, Yixiao Zhang

**Author notes:** These authors contributed equally (Xiaojie Wang, Zhongjian Bai, Ciara Wallis). To whom correspondence should be addressed: Dr. Yixiao Zhang, Dr. Ben Corry, Dr. Yadong Sun, Dr. Stephen Shears.

## Abstract

XPR1 is emerging as the only known inorganic phosphate (Pi) exporter in humans, critical for Pi homeostasis, with its activity stimulated by inositol pyrophosphate InsP8 and regulated by neuronal scaffold protein KIDINS220. Our structural studies reveal InsP8 specifically activates XPR1 in a stepwise manner, involving profound SPX domain movements. Each XPR1 subunit functions with four gating states, in which Pi permeates a constriction site via a “knock-kiss-kick” process. In contrast, KIDINS220 delicately stabilizes XPR1 in a closed conformation through multiple mechanisms, one of which involves trapping the XPR1 α1 helix—critical for InsP8 binding—within an interaction hub. InsP8 serves as a key to release KIDINS220’s restraint, reinforcing a “key-to-locks” mechanism to safeguard the stepwise activation. Additionally, our study provides direct structural insights into XPR1-associated neuronal disorders and highlights the evolutionary conservation and divergence among XPR1 orthologues, offering a comprehensive understanding of Pi homeostasis across species.

## Introduction

Inorganic phosphate (Pi) is an essential element for all living organisms and plays critical roles in various biological processes^1,2^. It is involved in the construction of genetic material, energy metabolism, signal transduction, phospholipid synthesis, hard tissue formation, and pH buffering ^3^. Therefore, the metabolism of Pi must be tightly regulated, as excess Pi in cells is toxic ^4^. While the transporters responsible for Pi uptake, including the SLC20 and SLC34 protein families, have been identified and extensively studied ^5–7^, XPR1 (SLC53A1) is emerging as the only protein known to mediate Pi efflux in human ^8–10^.

XPR1 was initially identified as a receptor for xenotropic and polytropic murine leukemia retroviruses ^11–13^. Later, it was shown to function as a Pi exporter when intracellular Pi levels increase ^8,9,14,15^. It is broadly expressed in different cell types and organs ^8^ and is well conserved across species. Impairment of XPR1 function leads to severe complications, including primary familial brain calcification (PFBC), hypophosphatemia, renal Fanconi syndrome, and tumorigenicity ^16–24^. Its orthologues—PHO1 in plants, SYG1 in yeast, and PXo in Drosophila—are all critical for Pi homeostasis ^25–27^, though little is known about the detailed molecular mechanism of this important protein family.

XPR1 contains three domains: an N-terminal SPX (SYG1/PHO/XPR1) domain, a transmembrane core domain, and a C-terminal EXS (ERD1/XPR1/SYG1) domain followed by a loop region. The SPX domain acts as a sensor for inositol polyphosphates (InsPP) in a wide range of organisms ^28^. When Pi concentration increases, PPIP5Ks and IP6Ks convert InsP6 to 1,5-InsP8 (referred to as InsP8), which serves as an intracellular Pi concentration indicator ^29^. InsP8 then activates XPR1 through the SPX domain, allowing Pi efflux via the EXS domain to reduce cellular Pi level^14,15,30–32^. In addition to InsP8, recent studies suggest that KIDINS220, a protein involved in neuronal signaling and development ^33–35^, can also form a complex with XPR1 and regulate Pi efflux ^23^. To elucidate the precise molecular mechanisms regulating XPR1, we determined cryo-EM structures of XPR1 in multiple functional states, including complexes with InsP8 and KIDINS220. Through a combination of structural, functional, and computational studies, we revealed that XPR1, a member of the SLC family, operates as a channel-like protein. It employs a “key-to-locks” mechanism to safeguard the stepwise activation and a “knock-kiss-kick” mechanism to facilitate Pi efflux.

## Results

### The architecture of XPR1 in closed state

We first overexpressed and purified full-length human XPR1 from 293F cells and performed cryo-EM studies (**Figure S1**, **Table S1**). From 2D averages and 3D cryo-EM maps, it is clear that XPR1 forms a homodimer **(Figures 1A-1C, Figures S1B-S1D)**. TM1 in the core domain forms the dimerization interface, where the lipid acyl chains also play a role in bridging the two subunits **(Figure 1C, Figures S1D-S1F)**. We observed several other lipids bound around XPR1, with one lipid buried inside the protein, sandwiched between TM2-5 and the membrane-contacting amphipathic helix IH1, likely playing a structural role **(Figure 1C, Figures S1D, S1G and S1H**). The EXS domain is located on the sides of XPR1 **(Figures 1B-1C)**, fully embedded in the membrane, with several charged and polar residues pointing toward the center, likely forming a pathway for Pi to pass through **(Figure 1D, Figure S1I)**.

**Fig. 1.**
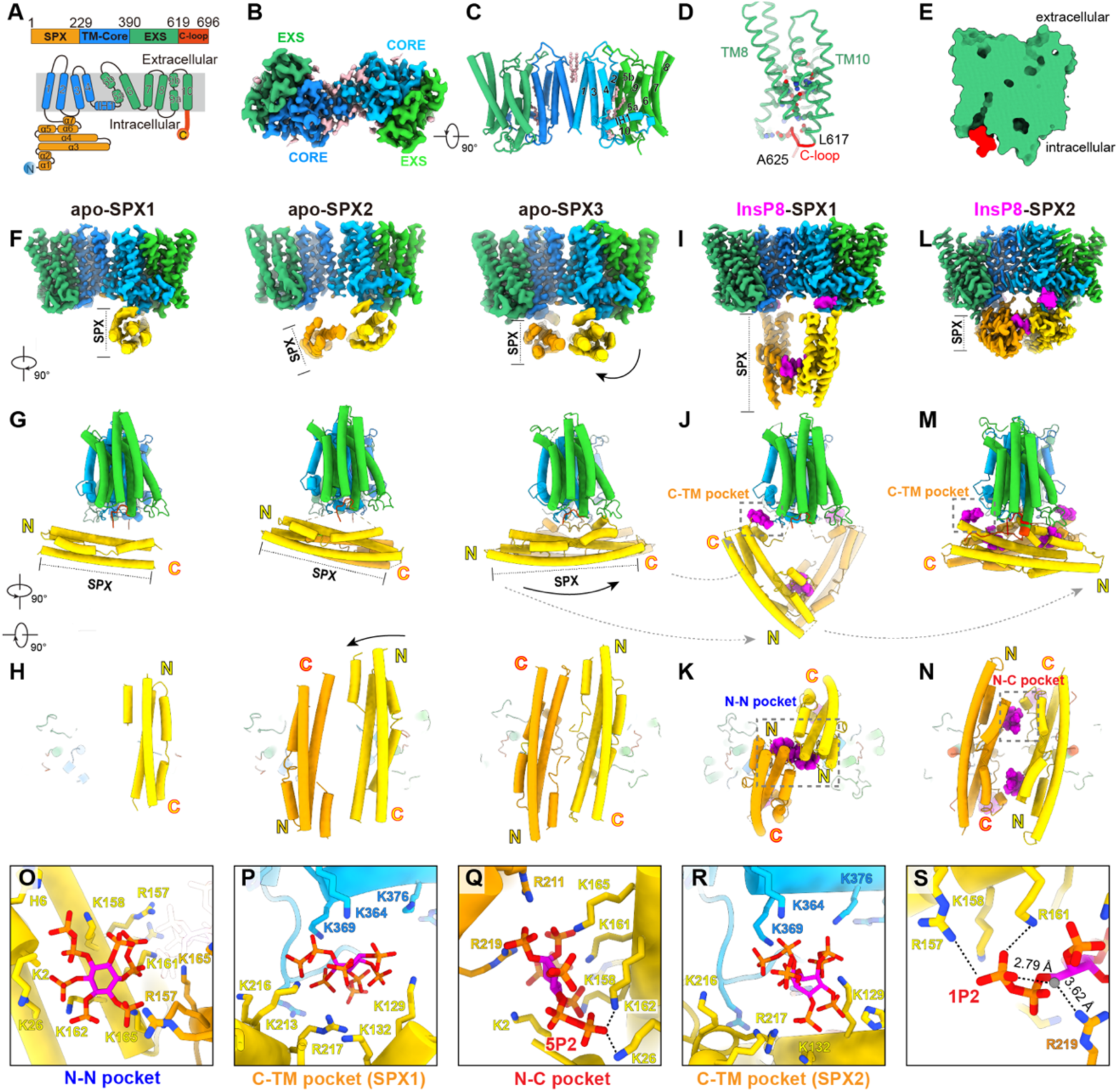
InsP8 binding stabilizes the dynamic SPX domain. **A**, A cartoon depiction of the domain organization in XPR1. **B**, The Cryo-EM map of XPR1 in the apo state is shown in the bottom view. The domains are colored according to (A). The lipid density is colored in pink. **C**, The model of XPR1 in the apo state, with selected lipids displayed as ball-and-stick in pink. **D**, The charged and polar residues in the EXS domain are displayed as ball-and-stick. The loop residues 617-625 covering the pore are colored as red. **E**, The clip surface of the XPR1 subunit shows that the permeation pathway is covered by the C-loop (red) on the intracellular side and tightly sealed on the extracellular side. **F**, The Cryo-EM maps of XPR1 in the apo-SPX1, apo-SPX2, and apo-SPX3 states. **G-H**, The models of XPR1 in the apo-SPX1, apo-SPX2, and apo-SPX3 states are shown in lateral (G) and bottom (H) views, with the movement of SPX domains indicated by arrows. **I-K**, The structure of XPR1 in the InsP8-SPX1 state is shown in side view (I), lateral view (J), and bottom view (K), with the pockets for InsP8 (magenta) binding outlined in dashed boxes. **L-N**, The structure of XPR1 in the InsP8-SPX2 state is shown in side view (L), lateral view (M), and bottom view (N), with the pockets for InsP8 (magenta) binding outlined in dashed boxes. **O-R**, Zoomed-in views of the InsP8 binding pockets. The N-N pocket (O) and C-TM pocket (P) in the InsP8-SPX1 state, and the N-C pocket (Q) and C-TM pocket (R) in the InsP8-SPX2 state are displayed. The positively charged residues are shown as sticks. **S**, Recognition of 1P2 in the N-C pocket in the InsP8-SPX2 state. A water molecule is displayed in gray.

The pathway is tightly sealed on the extracellular side, at the height of W573 **(Figure 1E, Figure S1J)**. In contrast, at the intracellular side, the TM helices form a wide opening **(Figure 1E)**. This opening contains several positively charged residues, consistent with Pi binding capacity **(Figure S1K)**. However, residues 619-625 form a loop that largely covers the pore, leaving the opening a narrow one **(Figure 1E, Figure S1J)**. The C-terminal residues after 625 are missing from the EM density. Our radius profile suggests that this loop significantly reduces the pore radius at the opening from 3.4 Å to 1.3 Å **(Figure S1J)**. Considering the covered opening on the intracellular side and the tightly sealed extracellular gate, we suggest that this structure represents a closed state of XPR1.

### The SPX domain is highly dynamic in apo states

Though the transmembrane core and EXS domains are clearly resolved, the intracellular SPX domain, which is key for InsP8 sensing, is invisible in our high-resolution cryo-EM structure, indicating it is highly dynamic. In fact, during our 2D and 3D classifications, in some classes, we were able to observe the SPX domain in different orientations (apo-SPX1,2,3) **(Figures 1F-1H, Figure S1L)**. In apo-SPX1, only one SPX domain is visible, oriented perpendicular to the transmembrane helices and lying below the membrane. In apo-SPX2, two SPX domains dimerized and asymmetrically floated below the transmembrane region, with only one SPX attached to the TM region. In apo-SPX3, two SPX domains lay below the TM region in a symmetric manner. Rotations and tilts of the SPX domains are observed between these three states, further indicating their dynamics **(Figures 1F-1H)**. We expected that the different SPX conformations might play a role in regulating the conformations of the TM region, and thus XPR1 activity. However, although the SPX domains adopt different conformations, the TM region in these structures remains nearly identical, all in the same closed state **(Figure S1M)**. This raises the question of how InsP8 binding can activate XPR1 through the SPX domain.

### The dynamic SPX domain is stabilized by InsP8 at different states

Next, we aimed to understand InsP8-dependent XPR1 activation by obtaining the structure of the XPR1-InsP8 complex. By adding chemically synthesized InsP8 ^36^ in the cryo-EM sample, we found the SPX domain become visible in most 2D and 3D classes, indicating SPX domain was stabilized with InsP8 **(Figure S2)**. Interestingly, we observed the SPX domain in two distinct conformations (XPR1-InsP8-SPX1 and XPR1-InsP8-SPX2) **(Figures 1I-1N)**.

In XPR1-InsP8-SPX1, the SPX undergoes a significant conformational change compared to its apo states. The two SPX domains tilt upwards, shifting from a membrane-laid orientation to an “arch-like” conformation **(Figures 1I-1K)**. Even more remarkably, the two SPX domains flip their orientation beneath the TM region. This means the SPX domains do not simply tilt up, but each elongated SPX domain rotates approximately 180 degrees along the axis perpendicular to the membrane, causing the N-terminal and C-terminal regions of the SPX domains to switch orientations **(Figures 1J-1K)**. In XPR1-InsP8-SPX2, the SPX domain lies below the membrane **(Figures 1L-1N)**. At first glance, it appears to be in a similar position as in XPR1-apo-SPX3; however, the domains are also flipped, with switched N-terminal and C-terminal orientations, and each SPX domain rotates approximately 80° along its elongated axis after the flipping **(Figures 1M-1N, Figure S2C)**. These remarkable conformational changes demonstrate the ability of InsP8 to regulate SPX domains.

### The specific recognition of InsP8 by SPX domain

In each of these two classes, we observed four InsP8 molecules in two binding pockets **(Figures 1O-1S)**. In XPR1-InsP8-SPX1, two InsP8 molecules bind at the tip region of the arch, where the N-terminal of SPX forms a highly positively charged pocket to accommodate InsP8 binding. Each InsP8 is recognized by residues from both SPX domains (referred to as N-N pocket) **(Figures 1K and 1O)**. The other two InsP8 molecules bind in positively charged pockets formed by the SPX C-terminal and the membrane-laying IH1 (referred to as C-TM pocket) **(Figures 1J and 1P)**. These two InsP8 molecules are also present in the same location in XPR1-InsP8-SPX2 **(Figures 1M and 1R)**, likely playing a role in stabilizing the dynamic SPX in the flipped orientation by anchoring the C-terminal of SPX to the TM region. In XPR1-InsP8-SPX2, the N– and C-terminal of SPX are also involved in InsP8 binding, but in a different manner compared to XPR1-InsP8-SPX1. Two InsP8 molecules bind in positively charged pockets formed by the N-terminal SPX of one subunit and the C-terminal SPX of another subunit (referred to as N-C pocket) **(Figures 1N and 1Q)**. These two InsP8 molecules effectively glue the two SPX domains together. The two additional phosphates (1P2, 5P2) in InsP8 are specifically recognized. 5P2 interacts with Lys26 and Lys162 **(Figure 1Q)**, while 1P2 interacts with R157, K158, R161 from the same subunit and R219 from the other subunit through water molecule **(Figure 1S)**. These N-C pockets do not exist if the two SPX domains simply flip 180 degrees, indicating that in this flipped state, the 80 ° rotation along the elongated SPX axis is induced and stabilized by these two InsP8 molecules. In short, the InsP8 in the C-TM pocket likely helps stabilize SPX in the flipped orientation, while the InsP8 in the N-N and N-C pockets helps stabilize SPX in the “arch-like” and “laid-rotated” states, respectively. These specific interactions support the role of InsP8 in regulating XPR1 activity.

### The opening of intracellular gate induced by InsP8

Next, we wanted to ask whether these InsP8-induced conformational changes can affect the conformation of the transmembrane region. In the “arch-like” XPR1-InsP8-SPX1 structure, although significant conformational changes were observed in the SPX domain, the TM region remained unchanged **(Figure S2D)**. However, in the XPR1-InsP8-SPX2 cryo-EM map, we found it could be a mixture of different conformations **(Figure S2E)**.

By performing careful focused 3D classification **(Figure S2B)**, we identified two distinct conformations on the intracellular side: one is the same as in the apo state, with a short loop largely covering the opening **(Figures 2A, 2C and 2E)**. In the other conformation, this loop straightens up and extends into a helix on TM10 **(Figures 2B, 2D and 2E)**. The following C-terminal residues, 626-632, which were unresolved in the apo state, are now visible and form extensive interactions with the side of the SPX domain **(Figures 2B and 2D)**. It is likely that the flipped and rotated SPX domain provides a binding platform for these flexible C-terminal residues, which, in turn, help release the cover loop, enlarging the pore radius from ∼1 Å to ∼3 Å at the intracellular gate **(Figure 2E)**.

**Fig. 2.**
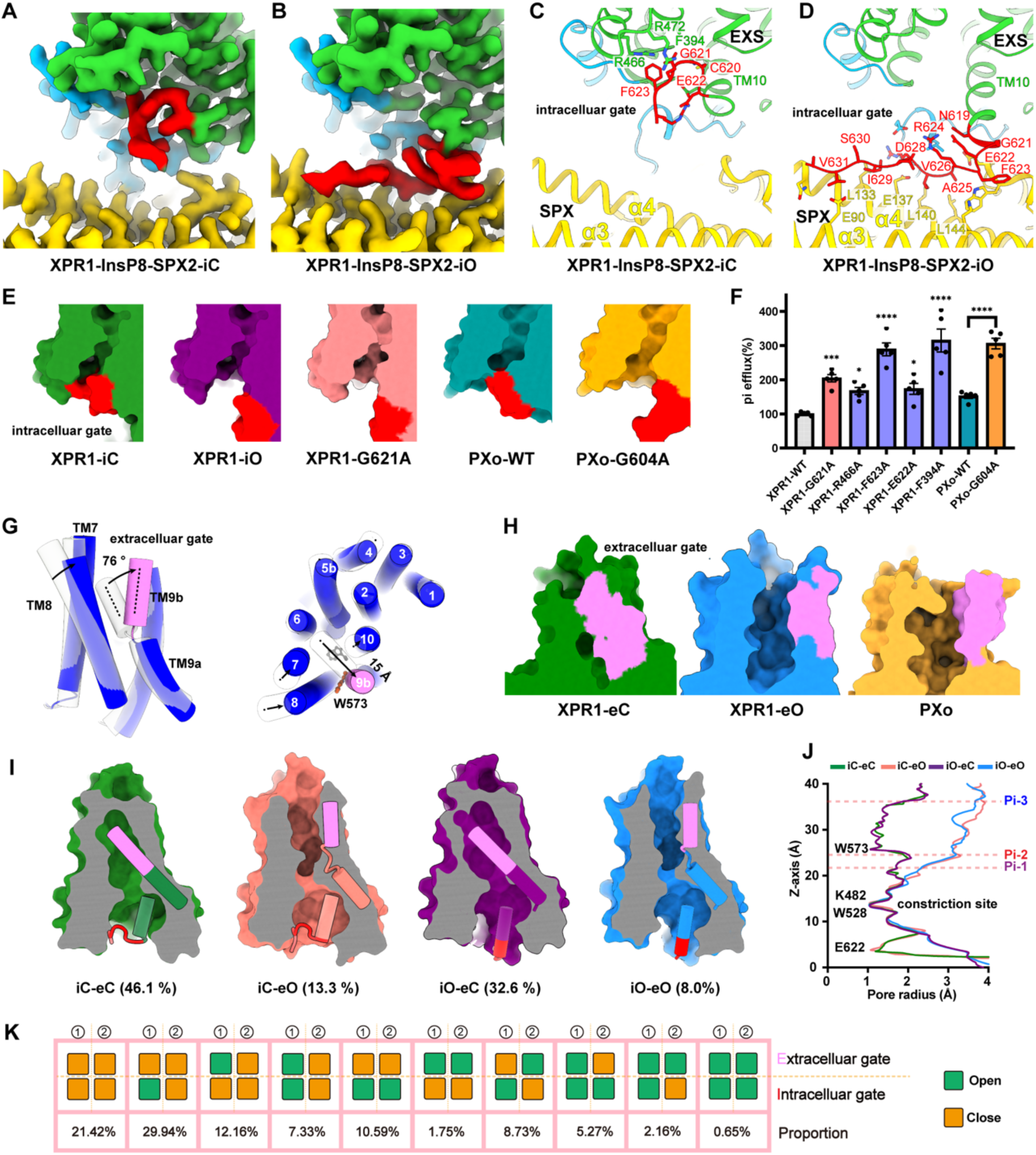
The opening of the intracellular and extracellular gates is uncoupled. **A-B**, Cryo-EM maps of XPR1 in the InsP8-SPX2-iC state (A) and InsP8-SPX2-iO state (B) show the conformational change of the intracellular gate. **C-D**, Models of XPR1 in the InsP8-SPX2-iC state (C) and InsP8-SPX2-iO state (D) illustrate the conformational change of the intracellular gate. The interacting residues are shown as sticks. **E**, Surface clips of XPR1-iC, XPR1-iO, XPR1-G621A, PXo-WT, and PXo-G604A depict the conformation of the intracellular gate. The C-loop residues are colored in red. **F**, Relative Pi efflux levels in cells expressing different XPR1 and PXo constructs. Data (n=5) from three independent assays are presented as mean ± SEM. A one-way ANOVA test was used; *P < 0.05, **P < 0.01, ***P < 0.001, and ****P < 0.0001. **G**, The conformation of the extracellular gate in closed (grey) and open (blue) states, with TM9b in the open state colored in violet. The arrows indicate the movement of the transmembrane helices from closed to open states. **H**, Surface clips of XPR1-eC, XPR1-eO, and PXo show the conformational change at the extracellular gate. The residues of TM9b are colored in violet. **I**, Surface clips of XPR1 in four different gating states, with the ratios of the four states labeled. TM9 and part of TM10 are shown as cylinders, with the C-loop colored in red and TM9b colored in violet. **J**, The pore radius along the permeation pathway of XPR1 in iC-eC, iO-eC, iC-eO, and iO-eO states, with the height of the bound Pi indicated by dashed lines. **K**, The cartoon illustrates the ratios of different gating states in the dimerized XPR1 protein. The squares denote the intracellular and extracellular gates, with the open gates represented in green and the closed gates in orange.

To provide functional evidence for the importance of this loop release, we first attempted to remove the cover loop and the following C-terminal residues; however, these proteins did not express well in our hands. We then aimed to screen mutants that could recapture this intracellular open conformation. We found that the G621A mutant in the cover loop results in a nearly identical open conformation at the intracellular gate **(Figure 2E, Figure S3A)**, and this mutant, which is not located inside the pore **(Figure 2C)**, significantly increased Pi efflux compared to WT XPR1 **(Figure 2F)**. Increased Pi efflux was also observed in other mutants of residues on the cover loop, such as E622A, F623A, and their interacting residues R466A and F394A **(Figures 2C and 2F)**. We further confirmed this observation using the Drosophila orthologue PXo, where the G604A mutant (corresponding to G621A in XPR1) straightened the cover loop and increased Pi efflux **(Figures 2E and 2F, Figures S3B and S3C)**. These data suggest that opening the intracellular gate of XPR1 and PXo can both facilitates Pi efflux.

### The opening of extracellular gate

At the extracellular side, we also observed two distinct conformations **(Figure 2G, Figure S2F)**: one is the same as in the apo state, with the extracellular gate tightly sealed. In the other, with a smaller particle population, TM9 forms a kink at residue W573, breaking into TM9a and TM9b **(Figures 2G and 2H)**. TM9b tilts outward by 76 °, and the tip residue T582 moves outward by 15 Å. The neighboring TM7, 8, 10 also exhibits subtle movement **(Figure 2G)**. The side chain of residue W573 near the kink flips away from the center, enlarging the pore radius from 1.1 Å to 3.1 Å, resulting in an open extracellular gate **(Figures 2G and 2H)**. This open conformation was also observed in the Drosophila PXo, in both the WT and the G604A cover loop mutant protein **(Figure 2H, Figures S3B and S3C)**, suggesting a conserved mechanism in Pi gating and permeation.

### The opening of the intracellular and extracellular gates is not coupled

Next, we wanted to understand if there’s coupling between intracellular and extracellular openings in XPR1, and if the two subunits need to work synchronously. By assigning the different states back to their original particles, we found that the XPR1 subunit can exist in four different states: iC-eC, iO-eC, iC-eO, and iO-eO (where lowercase ‘i’ and ‘e’ stand for intracellular and extracellular, and uppercase ‘C’ and ‘O’ stand for Closed and Open, respectively) **(Figures 2I and 2J, Figure S2G)**. From the subunit population, the fully closed iC-eC state has the highest ratio of 46.1%, whereas the fully open iO-eO state has the lowest ratio of 8% **(Figure 2I)**. Interestingly, we didn’t observe a significant difference in the ratios of extracellular opening in the intracellular gate closed (13.3/(46.1+13.3)=22.4%) and open (8/(32.6+8)=19.7%) state. These data suggest that extracellular opening does not seem to be affected or coupled with intracellular opening, and each gate can open independently **(Figure 2I)**. Moreover, at the particle level, we found that the two subunits also work independently, meaning that each subunit does not necessarily change symmetrically or adopt the same conformation in the dimerized particle **(Figure 2K)**. Since the conformational changes of the intracellular and extracellular gates do not appear to be coupled, and given the observation of the fully open iO-eO state, it is reasonable to suggest that XPR1 functions as a unique channel-like protein for Pi permeation.

### Pi recognition and conduction by XPR1

With the high-resolution structures of XPR1 in its activated states, we can analyze how Pi is recognized and permeated. To understand Pi recognition, we used the structure of the iO-eC state as an example for better local resolution, and it is also the state right before permeation. Three Pi ions were identified **(Figures 2J and 3A)**, with Pi-1 and Pi-2 locate between the heights of K482 and W573, while Pi-3 is positioned further towards the extracellular side. Several charged and polar residues are involved in Pi recognition, with the environment around Pi-1 appearing to be highly selective. The positively charged Arg459, Lys482, Arg570, Arg603, and Arg604 may play a role in attracting and holding Pi-1, while the negatively charged Glu398 and Glu529 balance the charge to maintain a suitable environment for Pi loading and release **(Figure 3B)**. The mutations of Arg459, Arg570, Arg604, Glu529, and Y483 at this site significantly reduced Pi efflux **(Figure 3E)**. Pi-2 is located near the residue W573 **(Figure 3C)**, which would flip out when TM9b tilts. Thus, Pi-2 is likely to be released immediately in the open conformation. The mutation of W573 at this site impairs Pi efflux, while S404 and T486 has a modest effect **(Figure 3E)**, supported by the fact that these two residues are relatively less conserved compared to W573 **(Figure S4A)**. Pi-3 is surrounded by Arg448, Arg273, and T580 further on the extracellular side **(Figure 3D)**, with the mutation of Arg448 having little effect **(Figure 3E)**, consistent with its location, which is far from the gating residues. Many of the pore-facing residues, especially those at the Pi-1 site, are highly conserved among XPR1 and its orthologs **(Figure S4A)**, suggesting their crucial role in Pi permeation.

**Fig. 3.**
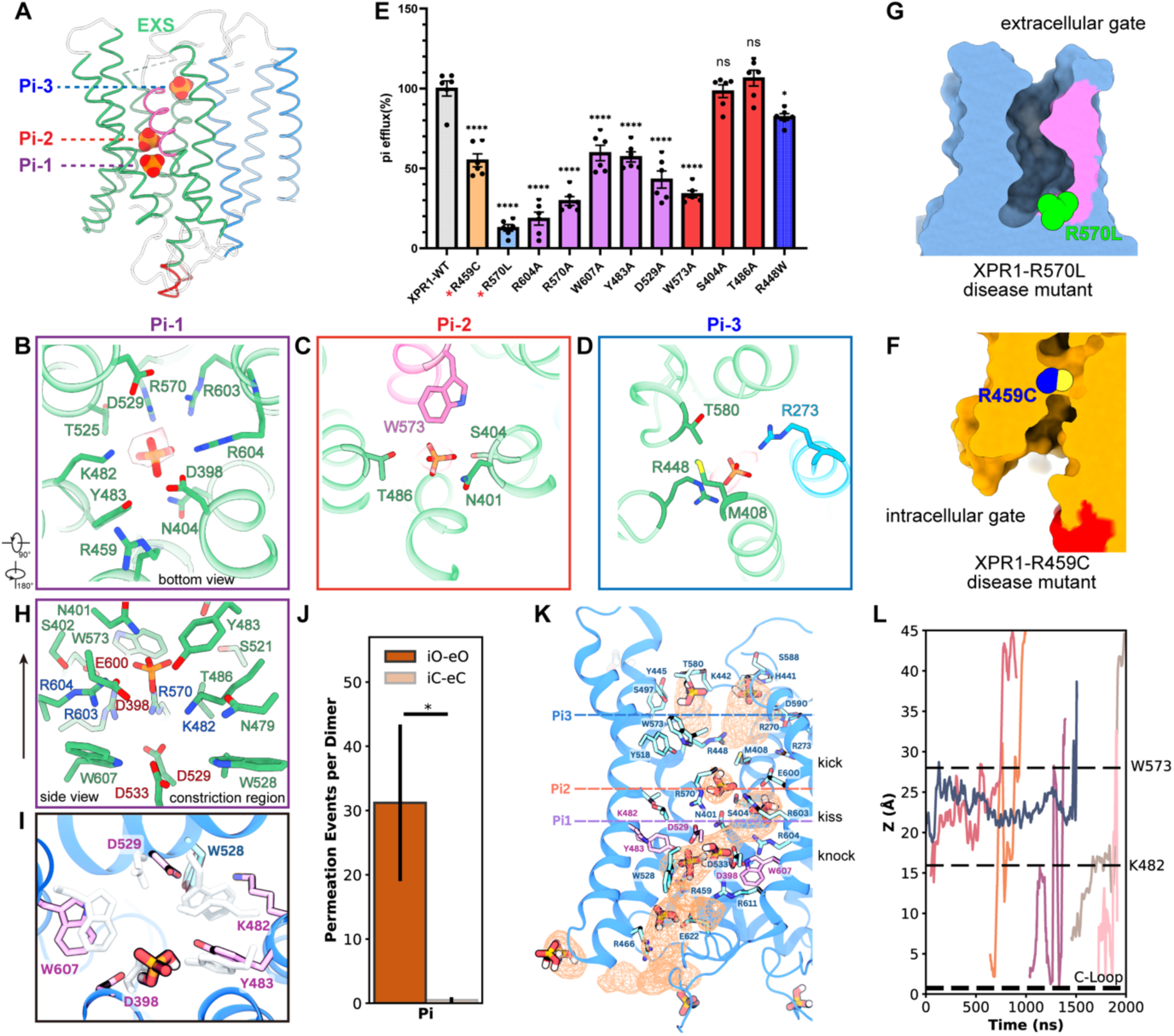
Pi recognition and the “knock-kiss-kick” model of Pi permeation. **A**, The positions of the bound Pi in the XPR1-iO-eC state. Pi1, Pi2, and Pi3 are displayed as spheres. **B-D**, The recognition of Pi1 (B), Pi2 (C), and Pi3 (D) by XPR1. The residues at the recognition site are displayed as sticks. **E**, Relative Pi efflux levels in cells expressing different XPR1 constructs. A red asterisk indicates the PFBC disease-related mutants. Data (n=6) from three independent assays are presented as mean ± SEM. A one-way ANOVA test was used; *P < 0.05, **P < 0.01, ***P < 0.001, and ****P < 0.0001. **F-G**, Surface clips of XPR1-R459C (F) and XPR1-R570L (G) show the open conformations of the intracellular and extracellular gates, respectively. The residues of R459C and R570L are displayed as spheres, while the C-loop and TM9b are colored in red and violet, respectively. **H**, The constriction region in the XPR1-iO-eC state. The relationship with panel (B) is displayed on the side of that panel. The arrow on the left indicates the direction of Pi permeation. **I**, Top-down view showing widening of the constriction formed by D398, K482, Y483, W528, D529, W607 observed in simulations of the iO-eO structure during a Pi permeation event. The starting positions of each residue in the iO-eO structure are shown in white and the position from the simulation in colors. **J**, Average total number of Pi permeation events (± SEM) observed at –400 mV for the iO-eO and iC-eC structures from simulations containing just the H₂PO₄⁻ species of phosphate. Statistical significance was determined using unpaired t-tests; *P < 0.05. **K**, Snapshot of a chain of Pi molecules along the transport pathway in the iO-eO structure. Regions commonly occupied by Pi are shown by the average volumetric density of Pi across the simulation (brown mesh, resolution 1 Å). **L**, Movement of representative permeating Pi ions through the transport pathway over time. The C-Loop, K482, and W573 positions are given as a reference. Each line indicates representative traces of Pi.

Among these pore residues, two positively charged residue mutants, R459C and R570L near Pi-1 site, were reported to cause primary familial brain calcification ^21,24^, a neurological disease attributed to the accumulation of cytosolic Pi and subsequent cerebral calcium-phosphate deposition. Unexpectedly, our structural studies show that R459C recaptured the intracellular opening state with the covered loop released, likely due to the reduction of positive charge and changes in the local environment, loosening the interaction with the covered loop **(Figure 3F, Figure S5A)**. Moreover, R570L, which is right below the TM9 kink residue, resembled the architecture of the extracellular opening state with TM9b tilted **(Figure 3G, Figure S5B)**. Though both mutants exhibit opening of the gating residues, they indeed impair Pi efflux in our and previous functional studies^21,37^ **(Figure 3E)**. This further emphasizes the critical roles of pore residues in Pi recognition and gating.

One important thing to note is that, although our structural studies show the opening of the intracellular and extracellular gates, a constriction site between W528 and K482 exists in all four gating states, including the fully open iO-eO state **(Figures 2J and 3H)**. This raises the question of how Pi passes through this narrow constriction site to entry the Pi-1 site. To answer this question, we performed MD simulation study **(Figures S5C-S5F)**. In the simulations, the side chains of many pore-lining residues are mobile. W528, and the acidic residues D529 and D398 at the constriction site can be pushed away by Pi **(Figure 3I)** allowing Pi permeation that does not occur in the iC-eC structure **(Figure 3J)**. In fact, this section of the pore widens on average during our simulations thus the constriction does not prevent Pi permeation **(Figures S5F)**. After the constriction site widens, a large number of Pi are observed entering the pore, attracted by the following positively charged residues R570, R603, and R604, forming a continuous chain across to the extracellular side of the protein **(Figure 3K)**. Pi tends to bind at this position longer than at others **(Figure 3L)**, suggesting that the layer of positively charged residues R570, R603, and R604 can attract and hold Pi **(Figures 3B, 3H and 3K)**. Interestingly, acidic and polar residues exist in the same region, such as D398, E600, Y483, N401, S402, and T486 **(Figures 3B, 3H and 3K)**; these residues may play a role in balancing the charge and providing a hydrophilic environment for Pi release. Based on the simulation data, we propose a “knock-kiss-kick” model for Pi permeation: Pi “knocks” the door open by pushing the mobile constriction residues, including acidic ones, widening the pore. Positively charged residues then “kiss” Pi where it dwells briefly. But acidic and polar residues prevent tight binding and “kick” the Pi toward the wider extracellular opening.

### The overall architecture of XPR1-KIDINS220 complex

Through the extensive structural studies described above, we learned the molecular details of how XPR1 is activated when in complex with InsP8. Recent studies suggest that XPR1 can also form a complex with KIDINS220 for proper localization and functioning ^23^. KIDINS220 is a protein preferentially expressed in the brain, involved in neuronal survival, maturation, and activity ^38^. The knockout of KIDINS220 resulted in reduced XPR1 surface expression, decreased Pi efflux, and cell death ^23^. KIDINS220 contains an N-terminal ankyrin repeats domain (AR), a four-helix transmembrane domain, a C-terminal SAM domain, and a PDZ domain **(Figure 4A)**. Deletion of the EXS domain of XPR1 abolishes its interaction with KIDINS220 ^23^, raising the question of whether KIDINS220 can interact with the EXS domain and regulate XPR1 activity. To verify their interaction, we co-expressed XPR1 with KIDINS220 (1-1076, removing the disordered C-terminal loop) and found that XPR1 can bring KIDINS220 from cytosol to the cell membrane **(Figure 4B)**, and the overexpressed proteins can co-elute in gel filtration **(Figure 4C)**, suggesting that these two proteins can form a complex in vivo and in vitro. Using these purified proteins, we were able to obtain a high-resolution structure of the XPR1-KIDINS220 complex **(Figures 4D and 4E, Figure S6)**.

**Fig. 4.**
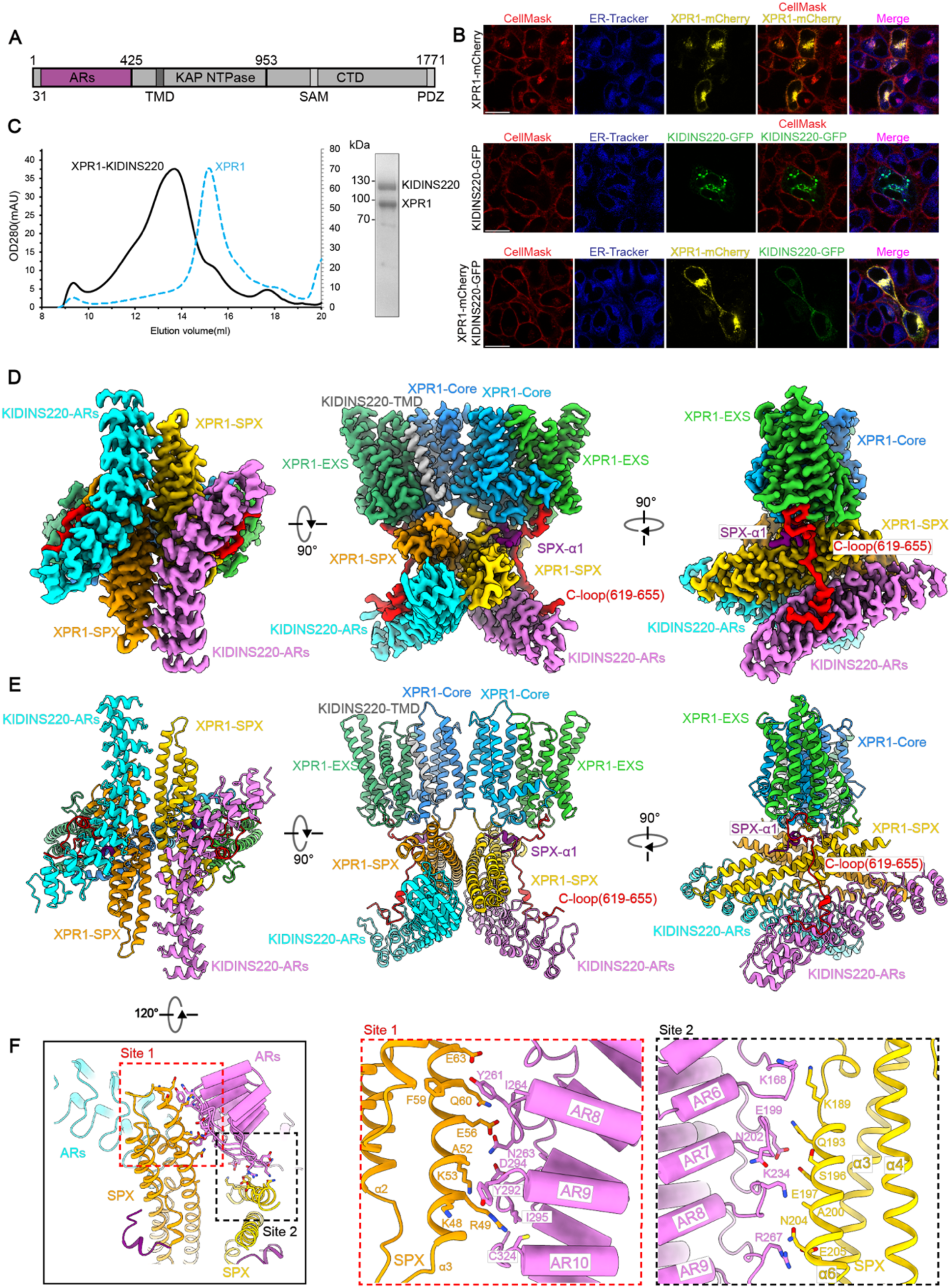
The overall architecture of the XPR1-KIDINS220 complex. **A**, Cartoon depiction of the domain organization of KIDINS220. **B**, XPR1 alters the localization of KIDINS220 from the cytosol to the cell membrane. CellMask (red) and ER-tracker (blue) were used to indicate the membrane and cytosol localization of XPR1 (yellow) and KIDINS220 (green) in HeLa cells (scale bar: 20 μm). **C**, Gel filtration profile of XPR1 alone (blue dashed line) and XPR1 in complex with KIDINS220 (black solid line). A left shift of the peak indicates the increased molecular weight of XPR1 in complex with KIDINS220. The Coomassie blue staining of the peak fraction of XPR1-KIDINS220 is displayed on the right. **D-E**, Cryo-EM map (D) and model (E) of the XPR1-KIDINS220 complex in bottom (left), side (middle), and lateral (right) views. **F**, The interaction between the KIDINS220 AR domain and the XPR1 SPX domain. Each AR domain binds with two SPX domains. The interacting residues are shown as sticks.

In this structure, XPR1 and KIDINS220 form a 2:2 complex with three major interaction regions **(Figure 4D and 4E)**: (1) two AR domains of KIDINS220 interact below the XPR1 SPX domains; (2) the XPR1 C-loop acts as a hook to dock the AR domains from the side; and (3) in a minor class, one TM helix from KIDINS220 interacts with the XPR1 EXS domain. The occupancy and the local resolution of this additional TM helix are not high enough for a reliable model building, so its identity was not assigned. Among these interaction regions, the AR-SPX interaction contributes to a large buried area of 1612 Å^2^. Each AR domain forms interactions with both SPX domains, specifically the connecting loops in AR8-10 interact with α3 in one SPX domain, and the connecting loops in AR6-9 interact with α6 in another SPX domain **(Figure 4F)**. These interactions may play a role in locking and stabilizing the dynamic SPX domains in this InsP8-free state.

### KIDINS220 stabilizes XPR1 in a closed state

Different from the unflipped, dynamic SPX domain observed in the InsP8-free apo state and the flipped SPX domain observed in the InsP8-binding states, in this structure, SPX is stabilized in a new conformation. Overall, it maintains an unflipped orientation like those in the apo states, but it undergoes an 100 ° rotation along the SPX elongated axis compared to apo-SPX3 **(Figure 5A)**. This conformational change causes the two SPX domains to form direct interactions through their α3 helices, leading to dimerization, which can further stabilize the conformation **(Figure 5A, Figure S6D)**.

**Fig. 5.**
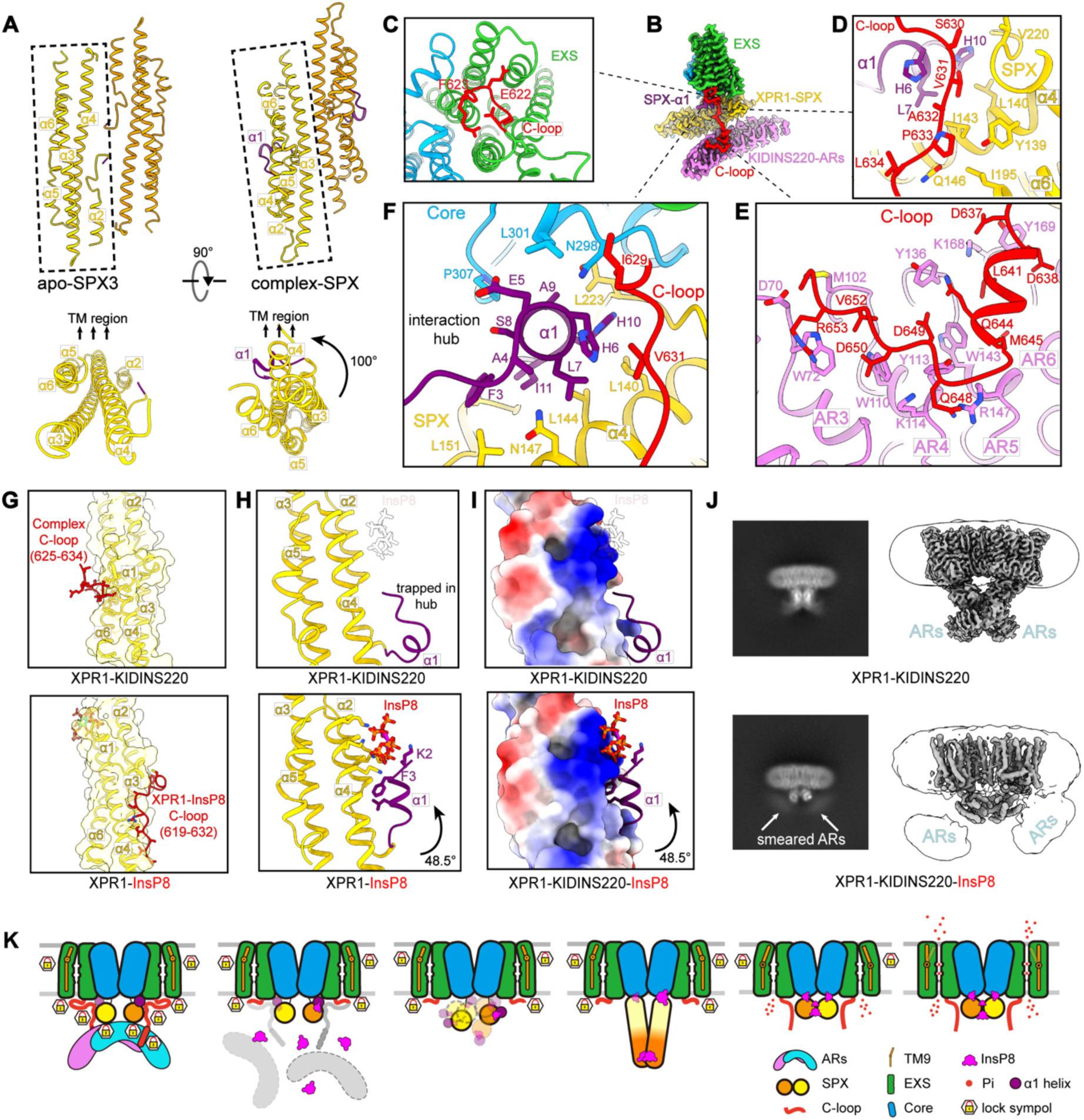
InsP8 acts as a key to release KIDINS220’s restraint. **A**, The conformational change of the SPX domain between the apo-SPX3 state (left) and the KIDINS220 bound state (right). The SPX domain rotates approximately 100° along the elongated axis upon KIDINS220 binding. **B**, The Cryo-EM map of the XPR1-KIDINS220 complex, the same as in Fig. 5D, indicates the position of the zoom-in views. **C**, The C-loop (red) covers the intracellular gate in the XPR1-KIDINS220 complex. **D**, The interaction between the XPR1 C-loop (red) and the SPX domain (yellow). The interacting residues are shown as sticks. **E**, The interaction between the XPR1 C-loop (red) and the KIDINS220 AR domain (violet). The interacting residues are shown as sticks. **F**, The SPX α1 helix forms an interaction hub. The SPX α1 helix (purple) interacts with C-loop residues (red), residues 298-307 on the loop connecting TM2 and TM3, residue L223 on the loop connecting the SPX and TM domains (blue), and residues on the SPX α4 helix (yellow). **G**, The XPR1 C-loop binds exclusively to different sides of the SPX domain (yellow) in the XPR1-KIDINS220 complex (upper panel) and the XPR1-InsP8 complex (lower panel). **H**, The conformational change of the SPX α1 helix between the XPR1-KIDINS220 complex (upper panel) and the XPR1-InsP8 complex. The SPX α1 helix tilts 48.5° upon InsP8 binding. In the upper panel, one InsP8 molecule is drawn transparently with a black outline to indicate its binding position on the SPX domain. **I**, The conformational change of the SPX α1 helix between the XPR1-KIDINS220 complex (upper panel) and the XPR1-KIDINS220-InsP8 complex. The SPX α1 helix tilts 48.5° upon InsP8 binding. The SPX domain is shown in surface representation and colored by electrostatic potential. **J**, Comparison of the 2D average and 3D map of the XPR1-KIDINS220 complex (left) and the XPR1-KIDINS220-InsP8 complex (right). Conformational changes are observed in the SPX and AR domains. The AR domain becomes smeared, and the SPX domain flips to the activated state with InsP8 added to the XPR1-KIDINS220 complex. **K**, A cartoon model showing the InsP8-dependent stepwise activation of XPR1.

The rotation of these unflipped SPX domains doesn’t result in the release of the C-terminal cover loop at the intracellular gate **(Figures 5B-5C)**, which aligns with our finding that the flipped SPX is crucial for XPR1 activation. Moreover, in this structure, the C-terminal residues (625-655) after the cover loop, which are missing in apo structures, are clearly seen and act like a hook to bridge the rotated SPX domain and the AR domain **(Figure 5B)**. Residues 630-634 interact with α1, α4, and α6 helices in the SPX domain **(Figure 5D)**, and residues 637-653 interact with AR3-6 of KIDINS220 at the concave side **(Figure 5E)**. These interactions likely enhance AR binding and stabilize SPX in this unflipped and rotated conformation. Interestingly, in our InsP8-bound activated structures, residues 619-632 interact with another surface in the SPX domain **(Figure 5G)**; thus, when KIDINS220 is present, these residues are occupied and not ready for the activated conformation.

Moreover, the rotation of the SPX domain brings the N-terminal α1 helix, which is involved in InsP8 recognition, to face the TM domain **(Figures 5A-5B)** and participates in forming an interaction hub **(Figure 5F)**. This short helix interacts with residues 298-307 on the loop connecting TM2-TM3, residue L223 on the loop connecting the TM domain and SPX domain, residues 629-631 on the C-loop, and residues 140-151 on the SPX α4 helix **(Figure 5F)**. All these interactions are new and specific to this AR-bound conformation.

Importantly, to accommodate these interactions, the α1 helix opens up and tilts approximately 48.5 degrees compared to the InsP8-bound state, resulting in a loosened pocket for InsP8 binding **(Figure 5H)**. The other way around, if the α1 helix is trapped in the interaction hub, it raises the question of whether InsP8 can still bind in this pocket, attract the α1 helix, and disrupt all these interactions, allowing for XPR1 activation. Our structural study demonstrated that by adding InsP8 to the XPR1-KIDINS220 complex, the closed state was released, and XPR1 transitioned to the activated state with a flipped SPX domain, in which KIDINS220 still binds but exhibits high dynamics and lacks defined structural features **(Figures 5I-5J, Figure S7A-S7B)**. Thus, InsP8, as a key, can disrupt these locking interactions in the XPR1-KIDINS220 complex and trigger activation.

In the TM region, the KIDINS220 TM helix interacts with TM9a in the EXS domain; although TM9b tilts toward this additional helix in the open state, they do not appear to conflict in either the closed or open state **(Figures S6E-S6F)**. Therefore, this structure does not provide sufficient evidence regarding whether this additional TM helix can regulate the EXS domain. However, the fact that the SPX domains are stabilized in the unflipped orientation, that the N-terminal α1 helix involved in InsP8 binding is altered and trapped on the TM domain, forming an interaction hub, and that the C-terminal residues important for intracellular gate opening are occupied by the SPX and AR domains all suggest that the binding of KIDINS220 may help stabilize XPR1 in a closed conformation when InsP8 is absent.

### The conformational cycle of XPR1 activation

Through our structural, functional, and computational studies, we have proposed a molecular pathway for the stepwise activation and Pi export of XPR1 **(Figure 5K)**. At low intracellular Pi concentrations, the SPX domain is either highly dynamic or stabilized by KIDINS220, with the latter stabilizing XPR1 in a closed conformation through an interaction hub formed with the SPX α1 helix. As intracellular Pi levels increase, the newly synthesized InsP8 binds to the SPX domain, attracting the α1 helix and inducing a significant conformational change. This change progresses from an arch-like intermediate state to a flipped and rotated state, creating a binding surface for the flexible C-terminal loop and causing the release of the covering loop at the intracellular gate. This release opens the intracellular gate, significantly enhancing phosphate access to the permeation pathway. The pore residues at the constriction site coordinate to facilitate Pi permeation, while the tilting of TM9b opens the extracellular gate, enabling Pi efflux **(Figure 5K, Video S1)**.

## Discussion

As one of the central elements in our homeostasis, phosphate homeostasis is much less understood in molecular detail compared to other ions such as Na⁺, K⁺, Ca²⁺, Mg²⁺, and Cl⁻. XPR1, the only reported Pi exporter, exhibits a unique feature: its activity is regulated by the Pi concentration indicator InsP8, suggesting that its activity needs to be precisely controlled and highly regulated. In this study, we revealed a “key-to-locks” mechanism for safeguarding the stepwise activation and Pi conduction of XPR1 **(Figure 5K)**. The “locks” include the α1 helix of XPR1, the C-terminal hook of XPR1, and the AR domain of KIDINS220, which lock the SPX domain in an unflipped state; the C-terminal loop that covers the intracellular gate; the constriction residues that narrow the permeation pathway; and TM9b, which closes the extracellular gate. InsP8 acts as the key to open these locks and facilitate Pi efflux. These mechanisms ensure that Pi is efficiently translocated only when phosphate levels are sufficiently high.

When the cellular Pi level is high, InsP6 is converted to InsP8, which serves as a Pi concentration indicator and the authentic activation ligand for XPR1. Thus, the two additional phosphates are critical for XPR1 substrate recognition and activation. A recent study reported the structure of the XPR1-InsP6 complex ^37^, which shows a very similar open conformation to our XPR1-InsP8-SPX2 structure. However, the two SPX domains in our InsP8 structure are closer together, due to the two additional phosphates, 1P2 and 5P2, in InsP8 **(Figures 6A-6B)**. This shorter distance may create a favorable binding geometry for the C-loop residues on the lateral side, potentially increasing the open probability of the intracellular gate. Indeed, in our XPR1-InsP8-SPX2 state, the open probability of the intracellular gate is 40.6% **(Figure 2I**, 32.6%+8%=40.6%), even in the presence of InsP8, indicating a relatively weak ability of the C-loop and the SPX domain to form interactions, which could play a regulatory role and has a higher demand for favorable binding geometry. Beyond shortening the distance between the two SPX domains for a potentially higher open probability, the “arch-like” state **(Figure 1I)**, which is unique in our InsP8 sample, may function as a “transit station”. This state may increase the likelihood of the flipping of the SPX domain to the active state, which is a significant and potentially challenging movement. Additionally, the higher binding affinity of InsP8 to the SPX domain ^28^ ^14^ and its potential greater capacity to unlock the KIDINS220 complex may also contribute to its higher selectivity over InsP6 and InsP7 for XPR1 activation, which could be further investigated.

**Fig. 6.**
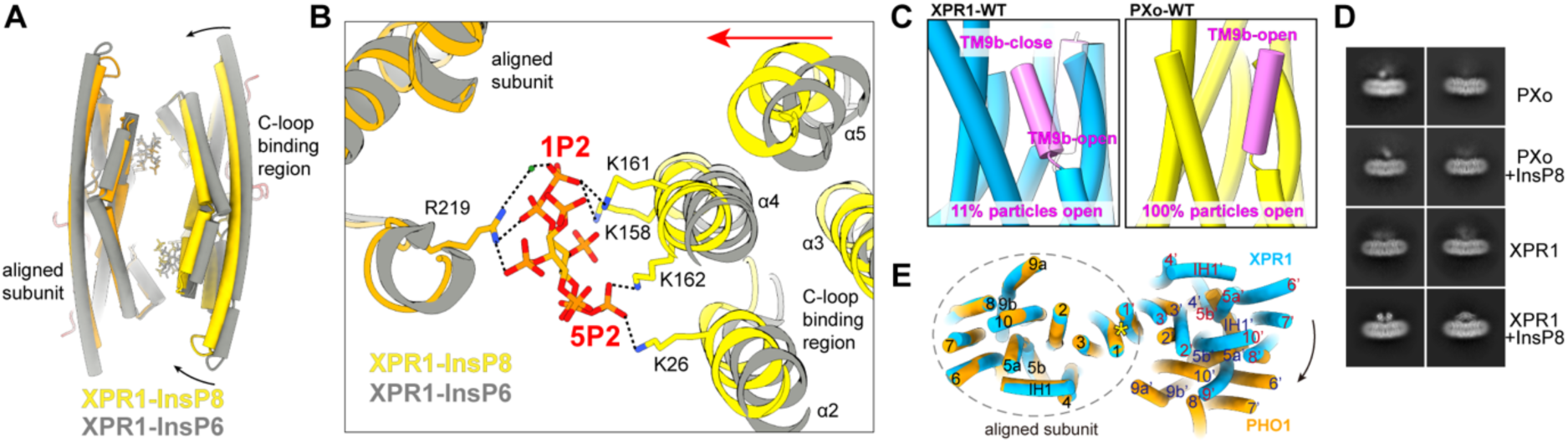
The two additional phosphates on InsP8 tighten the SPX domain and the structural difference among XPR1 orthologues. **A**, Structural comparison of XPR1-InsP6 (grey) and XPR1-InsP8 (orange), highlighting a closer SPX conformation in XPR1-InsP8. **B**, The two additional phosphates on InsP8 tighten the SPX domain. The residues involved in recognizing 1P2 and 5P2 are shown as sticks. A water molecular is colored in green. **C**, The TM9b (violet) of PXo-WT (right) has a larger open probability (100%) than XPR-WT (left, 11%). **D**, Comparison of the 2D averages of PXo and XPR1, with and without the addition of InsP8. **E**, PHO1 exhibits an asymmetric configuration. Structural comparison of XPR1 (blue) and PHO1 (orange), with the left subunit used for alignment. The C2 symmetry in XPR1 is indicated by a yellow asterisk.

Most transporters utilize an alternate accessing mechanism such as rocker-switch, rocker-bundle, and elevator models for substrate translocation^39^. However, our study suggests that the SLC family protein XPR1, with a novel SLC fold, acts as a channel-like protein, in which the opening of intracellular and extracellular gates generates a wide permeation pathway with a constriction site retained. Pi permeates through this constriction site using a unique ‘knock-kiss-kick’ mechanism. This working mechanism adds a novel model for substrate translocation across the membrane for the SLC protein family.

Phosphate homeostasis is critical for all species. Mammalian XPR1 shares 30.7% sequence similarity with PHO1 in plants and 54.4% with PXo in Drosophila. PHO1 is essential for Pi transport from root cells into the xylem, which is crucial for plant growth and development ^25,30^. Recent study shows that PXo is involved in a phosphate-sensing organelle that regulates cytosolic Pi levels ^27^. Our structural studies demonstrated that XPR1, PXo and PHO1 share a conserved architecture **(Figures S1D, S3C, S7C and S7D)**, and our mechanistic understanding of XPR1 sheds light on the working mechanisms among these orthologues. However, there are three major differences we observed among these three proteins: 1) The extracellular gate of PXo is consistently open with TM9b tilted; we did not observe a closed extracellular gate in the 3D classification **(Figure 6C)**. 2) The addition of InsP6 or InsP8 to the PXo sample does not stabilize the SPX domain or induce activation **(Figure 6D)**. In fact, there is currently no evidence that PXo can be activated by InsP6 or InsP8. Therefore, PXo may be regulated by other molecules, making it an intriguing target for future research into its activation mechanisms. 3) The dimerized PHO1 doesn’t exhibit C2 symmetry; its symmetry is broken at the interface between TM1 and TM3 on one of the subunits **(Figure 6E)**. Whether this asymmetry is related to its conformational cycle is unclear. These similarities and differences among the three proteins highlight both evolutionary conservation and divergence, opening up more intriguing questions for future studies.

Our study provides important implications for XPR1-related neurodisorders and tumorigenesis. Several familial PFBC mutations in XPR1 have been reported, and our structural analysis shows that most of these mutations are located in regions critical for XPR1 activation and gating, such as R459C, R570L, N619D and R604Q **(Figure S7E)**. Recent studies have found that the polarized distribution of SLC20 and XPR1 in astrocytes is crucial for brain phosphate homeostasis ^40^, further supporting the emerging roles of XPR1 in brain function. In line with this, KIDINS220 interacts with several neurotrophic receptors and is essential for neuronal survival, differentiation, and synaptic plasticity^38^. Aberrant expression of KIDINS220 is linked to a range of neuropsychiatric disorders and neurodegenerative diseases, including Alzheimer’s disease^41^. Previous studies suggest that the gene expression of KIDINS220 and XPR1 is highly correlated across various tissues, indicating their potential co-function ^23^. Our study provides clear evidence that KIDINS220 can form a complex with XPR1, and our data also suggest XPR1 can alter KIDINS220 localization to the plasma membrane. Thus, it would be both interesting and important to further investigate whether XPR1 can regulate the neuronal function of KIDINS220, and to understand the role of the XPR1-KIDINS220 complex in neuronal and other systems, as well as in pathological processes such as neurodegenerative diseases and cancers. Notably, some PFBC disease mutations are located at the XPR1-KIDINS220 interaction interface, including L140, which plays an important role in forming the α1 interaction hub **(Figures 5D and 5F, Figure S7E)**, and V652, which interacts with the KIDINS220 ARs domain **(Figure 5E, Figure S7E)**. This provides support for the critical role of the XPR1-KIDINS220 complex in Pi homeostasis. As KIDINS220 is required for XPR1 surface expression and Pi efflux in OVISE cells ^23^, the exact role of KIDINS220 in stabilizing the closed conformation would be intriguing for future studies, as it may influence the threshold for XPR1 activation or may function during trafficking.

Compared to other recent structural studies of XPR1 ^37,42–44^, this study is the only one to utilize the authentic substrate InsP8 and to obtain the XPR1-KIDINS220 complex, providing us with a unique opportunity to reveal several key and novel findings: 1) multiple locks that maintain XPR1 in a closed conformation; 2) the specific recognition of InsP8 by XPR1, acting as a key for these locks; 3) six distinct states of the SPX domain that enable a stepwise activation process; 4) the uncoupled opening of the intracellular and extracellular gates; 5) a unique “knock-kiss-kick” mechanism that facilitates Pi passage through the constriction site in this channel-like protein; and 6) Structural insights into disease-related mutations, rationally designed gain-of-function mutations, and XPR1 orthologues. By integrating careful image processing and MD simulations, this study elucidates the full conformational cycle of XPR1 activation and gating, with important implications for Pi homeostasis and inositol phosphate signaling, the gating mechanisms of transporters and ion channels, and the pathogenesis of neuronal diseases and cancers.

## Methods

### Cell culture

*Spodoptera Frugiperda9* (Sf9, ATCC (American Type Culture Collection)) cells were cultured in Sf-900™ medium in shake flasks at 26°C to 28°C on an orbital shaker, protected from light. HEK293F cells (ATCC), deficient in *N-acetylglucosaminyl transferase I* (GntI^−^), were grown in SMM 293-TII Expression Medium (Sino Biological) in a Herocell C1 orbital shaker incubator (Radobio) set to 37°C, 110 rpm, and 8% CO₂. HEK293T (ATCC) and HeLa (ATCC) cells were cultured in Dulbecco’s Modified Eagle Medium (DMEM) supplemented with penicillin-streptomycin and 10% fetal bovine serum (Gibco).

### Plasmid preparation

The genes encoding full-length Human XPR1 (UniProt accession code: Q9UBH6), KIDINS220 (UniProt accession code: Q9ULH0), PXo (UniProt accession code: Q9VRR2) and PHO1 (UniProt accession code: Q8S403) were cloned into the pEG Bacmam vector ^45^, which includes a precision protease cleavage site, a C-terminal enhanced green fluorescent protein (GFP) tag, and a Strep-tag II at the C-terminus for purification and detection. Additionally, Plasmid expressing XPR1 fused to a C-terminal FLAG-tag was constructed based on the pEG Bacmam vector. Specific mutations in XPR1 and PXo were introduced using PCR-based site-directed mutagenesis. Plasmids encoding XPR1 and KIDINS220 for cell imaging were constructed based on the pEG Bacmam vector, with C-terminal-mCherry and C-terminal-EGFP tags, respectively.

### Protein expression and purification

For large-scale preparation of XPR1, PXo and PHO1, the baculovirus expression system was employed to overexpress target proteins in HEK293F (GntI^−^) cells. Briefly, pEG Bacmam plasmids encoding target proteins were transformed into DH10Bac cells (Weidibio) to generate bacmids. These bacmids were then transfected into Sf9 cells with Fugene HD (Promega) to produce recombinant baculovirus. After two rounds of baculovirus infection, the resulting P3 viruses were collected and used for protein expression. When HEK293F cells reached a density of 2.5 × 10⁶ cells/mL, P3 baculovirus was added at a 1:20 volume ratio. For the overexpression of XPR1-KIDINS220 complex, cells were transfected with plasmids encoding XPR1-FLAG and KIDINS220(1-1076)-3C-GFP using PEI (PolySciences). After 16 hours at 37°C, 10 mM sodium butyrate was added to boost expression at 30°C. Cells were harvested by centrifugation at 4,000g after 48 hours.

For the purification of XPR1, cells were lysed in a buffer containing 50 mM Hepes, pH 7.4, 300 mM NaCl, 1 mM DTT, 2 μg/ml DNase I and protease inhibitors (aprotinin, leupeptin, pepstatin, trypsin, benzamidine, PMSF), along with 1% n-dodecyl-β-d-maltoside (DDM) and 0.1% cholesteryl hemisuccinate (CHS). After a 2-hour solubilization, the lysate was centrifuged at 30,000g, and the supernatant was incubated with anti-GFP beads for 2 hours at 4°C. Beads were then washed with 20 column volumes of buffer (20 mM Hepes, pH 7.4, 150 mM NaCl, 0.05% DDM:0.005% CHS, 1 mM DTT) before overnight 3C protease treatment to cleave the GFP tag. Following cleavage, the flowthrough was concentrated and loaded onto a Superose 6 Increase column (Cytiva), pre-equilibrated with buffer (20 mM Hepes, pH 7.4, 150 mM NaCl, 0.05% GDN, 1 mM DTT). Target fractions were collected and concentrated for cryo-EM analysis. The same protocol was used for the purification of XPR1-G621A, XPR1-R459C, XPR1-R570L, PXo-G604A, PXo and XPR1-KIDINS220 complex. All steps were performed on ice or at 4°C.

### Cryo-EM sample preparation and data acquisition

The homogeneity of all samples was initially assessed by negative staining with 0.2% (w/v) uranyl acetate on a Tecnai T12 Transmission Electron Microscope (Thermo Fisher Scientific). For cryo-EM grid preparation of XPR1-apo, XPR1-G621A, XPR1-R459C, XPR1-R570L, PXo, PXo-G604A and PHO1, samples were incubated with 50 mM NaH_2_PO_4_ for 30 minutes prior to grid preparation. For XPR1-InsP8 cryo-EM grids, samples were incubated with 50 mM NaH_2_PO_4_ and 1 mM chemically synthesized InsP8^36^ for 30 minutes. For XPR1-KIDINS220-InsP8 cryo-EM grids, 1 mM InsP8 was added before grid preparation. All samples were applied to glow-discharged 200 mesh 2/1 Au grids (Quantifoil).

Cryo-EM data were collected using a 300 kV Titan Krios G4 microscope (Thermo Fisher Scientific) equipped with a Biocontinuum K3 Direct Electron Detector and a GIF filter (Gatan). Movie stacks were automatically acquired using EPU v3 (Thermo Fisher Scientific) at a magnification of 81,000×, with a pixel size of 1.055 Å. The dose rate was set to 25 counts per pixel per second, and movie stacks were fractionated into 40 frames with a total exposure time of 2.2 seconds. The defocus range was set between −1.4 and −2.4 μm.

### Cryo-EM image processing

For the XPR1-apo, XPR1-InsP8, XPR1-G621A, XPR1-R459C, XPR1-R570L, PXo, and PXo-G604A and PHO1 datasets, the collected movie stacks were gain-normalized, dose-weighted and motion-corrected in Motioncorr2 ^46^. CTF parameters were determined with CTFFIND4 ^47^. Particles were automatically picked with Gautomatch (https://www2.mrc-lmb.cam.ac.uk/download/gautomatch-053/), extracted from the micrographs and subjected to 2D classification in RELION-3.1 ^48^. Particles from 2D classes showing secondary structure features were selected for further 2D classification in CryoSPARC ^49^. Particles displaying clear secondary structure features were selected and subjected to multi-rounds of Ab-initio Reconstruction or Heterogeneous Refinement to determine different conformations while poor-quality particles were excluded.

For XPR1-apo dataset, the particles without a clear SPX domain underwent additional rounds of Ab-initio Reconstruction to remove the poor particle. The remaining particles were combined and refined using non-uniform refinement in CryoSPARC, resulting in high-resolution maps. Particles with clear SPX domain were subjected to several Ab-initio Reconstruction to resolve the different conformations of XPR1. The results revealed that the particles mainly clustered into three subclasses, with the SPX domain showing high heterogeneity. These subclasses were then individually refined using non-uniform refinement.

For XPR1-InsP8 dataset, Ab-initio Reconstruction reveal that the SPX domain exhibited two distinct conformations. The SPX domain displayed an “arch-like” conformation, referred to as InsP8-SPX1, and a flipped conformation referred to as InsP8-SPX2. After several rounds of Ab-initio Reconstruction, particles from InsP8-SPX1 class were combined and subjected to the non-uniform refinement. Particles of InsP8-SPX2 were subjected to non-uniform refinement, resulting in high-resolution maps that identified a mixture at the C-loop and extracellular gate site. To investigate the different states at the intracellular and extracellular gate, particles in InsP8-SPX2 class underwent symmetry expansion (C2), followed by local refinement with a mask applied to one monomer. Particles after local refinement were subjected to focused 3D classifications in CyroSPARC, specifically targeting the intracellular gate and extracellular gate individually. Subsequently, particles in different states iO (intracellular gate Open), iC (intracellular gate Close), eO(extracellular gate Open), eC(extracellular gate Close) were subjected to local refinement to obtain high-resolution maps. To determine the complete conformational states of the entire subunit, particle intersections in CryoSPARC were used to isolate overlapping particles. These particles were then subjected to local refinement, resulting in the identification of four distinct states: iC-eC, iO-eO, iO-eC, and iC-eO.

For the PXo and PXo-G604A, XPR1-G621A, XPR1-R459C and PHO1 datasets, selected particles in 2D classification job were subjected to multiple runs of Ab-initio Reconstruction to clean up poor-quality particles. Particles exhibiting clear 3D features were then subjected to non-uniform refinement. In the PXo-G604A dataset procedure, CryoSieve ^50^ was used to eliminate unnecessary particles from final particle stack resulting a higher resolution density maps in the final refinement.

For the XPR1-KIDINS220 complex and XPR1-KIDINS220-InsP8 datasets, collected movie stacks were imported into CryoSPARC, and gain-normalized, dose-weighted, and motion-corrected using Patch Motion Corr. CTF parameters were determined with Patch CTF. In the XPR1-KIDINS220 complex datasets, particles were picked using Topaz picking ^51^ and extracted from selected micrographs with binning factor of 2. The particles underwent 2D classification, and those particles from 2D class averages with clear secondary features were selected for Ab-initio Reconstruction and Heterogeneous Refinement to remove poor-quality particles. The best particles from heterogeneous refinement were further processed with non-uniform refinement. Subsequently, local refinement imposed with a mask covering KIDINS220 and SPX was conducted to improve the density of KIDINS220. For model building, maps from non-uniform refinement and local refinement were combined in UCSF Chimera and post-processed with EMReady_V2. For the additional TM helix, particles from non-uniform refinement were subjected to symmetry expansion, followed by local refinement using a soft mask covering a monomer of the XPR1-KIDINS220 complex. Subsequently, 3D classification focused on additional TM helix and part of XPR1-EXS was performed. Classes containing additional TM helix were selected for local refinement with a soft mask covering the TM domain of XPR1 and tbe additional TM helix.

For the XPR1-KIDINS220-InsP8 dataset, particles were picked using template picking and extracted from selected micrographs with a binning factor of 2. All extracted particles were subjected to 2D classification, and classes with clear secondary structure were selected for Ab-initio Reconstruction. Particles from 3D classes with clear structural features were re-extracted with a binning factor of 1 and refined using non-uniform refinement.

Fourier shell correlation curves (FSC=0.143) were calculated in CryoSPARC. For additional details regarding the number of micrographs, particle counts, and the processing workflow, please refer to the Supplementary figures and tables.

### Model building and pore radius plot

EMReady_V2 ^52^ was used to improve the interpretability of cryo-EM maps for the model building. The AlphaFold2-predicted ^53^ structures of human XPR1, PXo, PHO1 and KIDINS220 were fitted into the cryo-EM density map using UCSF Chimera and manually adjusted in Coot ^54^. The models were then optimized through iterative cycles of real-space refinement in PHENIX ^55^, alone with manual adjustments in Coot. The model-building statistics are summarized in Table S1. Pore radius calculations were performed using the HOLE program ^56^, while the phosphate permeation pathway was visualized in VMD ^57^. Plots were generated using GraphPad Prism 9, and the structural figures were prepared in UCSF Chimera ^58^ and UCSF ChimeraX ^59^.

### Cell based Pi efflux assay

The Pi efflux assay was conducted according to established methods^20^. HEK293T cells were seeded onto poly-D-lysine-coated 48-well plates (BIOFIL). The following day, the cells were transfected with plasmids encoding XPR1, PXo, and their mutations using PEI (PolySciences). After 24 hours, the cells were washed three times with phosphate-free DMEM (Gibco) and incubated for 1 hour. Supernatants (80 µl) from each well were transferred to a 384-well plate and mixed with 20 µl of Malachite Green reagent (Sigma) for 30 minutes and absorbance at 620 nm was measured using an EnSight Multimode Plate Reader (PerkinElmer). To evaluate expression levels of XPR1, PXo, and mutations, western blot analysis was performed. Band densities were quantified using ImageJ, and ratios of band densities were used to calculate relative expression levels of XPR1, PXo, and mutations. The final Pi efflux data were normalized to the relative expression levels and analyzed in GraphPad Prism 9.

### Western Blot assay

The cells from the Pi efflux assay were lysed in RIPA buffer, followed by sonication for 20 seconds and mixing with 5× SDS loading buffer. The samples were run on 10% SDS polyacrylamide gels at a constant voltage of 150V for 70 to 80 minutes until the dye moved out of the gels. Subsequently, the gels were assembled with activated PVDF membranes into the electroblotting cassettes and run in Tris-Glycine transfer buffer at 110V for 70 minutes (with the current under 0.4 A). The blotted membranes were washed 3 times with TBST buffer and blocked with freshly prepared 5% non-fat milk (Sangon Biotech) for 30 minutes. After blocking, the membranes were washed 3 times and incubated overnight at 4℃ with primary antibodies against GFP and GAPDH (Proteintech), diluted in Primary Antibody Dilution Buffer (Beyotime) at 1:1000 and 1:2000, respectively. The membranes were then washed three times for 15 minutes each with TBST and incubated with the HRP-conjugated secondary antibody, diluted in 5% non-fat milk TBST buffer at 1:2000 for 1 h at room temperature. After incubation, the membranes were washed for 15 minutes with TBST and treated with BeyoECL Plus (Beyotime) for 1 to 5 minutes. The blots were detected using chemiluminescence imaging system (Fusion, VILBER) and the images were analyzed by ImageJ.

### Cell image of colocalization

HeLa cells were seeded onto the 4-Chamber Glass Bottom Dish (CellVis) and were transfected with plasmids encoding XPR1-mCherry and KIDINS220-GFP using PEI 24 hours after seeding. After 48 hours, the cells were washed there times with HBSS (Gibco) supplemented with 1.8 mM CaCl_2_. The cells were subsequently incubated with freshly prepared staining solution containing CellMask™ Deep Red (Invitrogen) and ER-Tracker (YEASEN) for 10 to 15 minutes. After removing the staining solution, the cells were washed three times and imaged using a Leica TCS SP8 confocal microscope.

### Molecular dynamics

The closed and open cryo-EM dimeric structures of XPR1 were prepared for molecular dynamics (MD) simulations using the CHARMM-GUI Membrane Builder server ^60^. Only the transmembrane core and EXS domains (residues 230 to 625) were included to reduce system size. N– and C-terminal residues were acetylated and methylamidated, respectively. A disulfide bond was included between C415 and C440 in each subunit. Structures were inserted into a 16 × 16 nm^2^ model membrane composed of pure dipalmitoylphosphatidylcholine (DOPC) lipids. The position and orientation of XPR1 within the membrane was determined using the Predictions of Proteins in Membranes (PPM v2.0) server ^61^, with the main-axis of the protein aligned along the simulation box’s Z-axis. Systems were solvated with 0.15 M NaCl solution. The structural lipid in each monomer between TM2, TM4, and TM5 was added and modelled as DOPC lipid in coot 0.9.7 ^54^ and furthered refined in real-space using Phenix 1.21.1. ^55^ The overall system dimensions were approximately 16 × 16 × 12 nm^3^ and consisted of ∼300,000 atoms, including ∼730 DOPC lipids, ∼180 Na^+^ ions, ∼150 Cl^-^ ions, and 65,000 water molecules **(Fig S5C)**.

At cytosolic pH phosphate can exist as both hydrogenphosphate (HPO_4_^2-^) and dihydrogenphosphate (H_2_PO_4_^-^). To determine which species is more likely to pass through XPR1, simulations are first conducted in the open pore structure under a –400mV voltage with a mixture of both species present. While both can enter the cavity, currents of H_2_PO_4_^-^ are much larger than that of HPO_4_^2-^ **(Figures S5C-S5F)**, this species is the focus and is referred to as Pi for simplicity.

As the XPR1 transport pathway is lined by several protonatable residues, the optimal protonation states of all aspartate, glutamate, and histidine residues were predicted using continuous constant pH MD simulations (CpHMD) using a specialized version of GROMACS (v.2021) with constant pH MD module implemented^62^. For CpHMD simulations, the modified CHARMM36 forcefield ^63^ was used to parametrize interactions of the system. λ-dynamics based constant pH MD ^62^ was applied to interpolate the hamiltonians of the protonation states of titratable residues. Multisite representation was used for histidine residues. Steepest descent energy minimization was conducted followed by six sequential steps of equilibration (112 ns in total) with a gradual decrease in the restraining force applied to protein backbone and sidechain atoms. Protein, lipids, and ion-water groups including buffer particles were treated independently to increase accuracy. 200 buffer ions were added into system by replacing water molecules for maintaining the charge neutral condition during simulation, the minimum distance between protein/lipid atoms and buffer ions was 1.0 nm, and those buffer ions were fixed in space during titration calculation. The pH range of simulation was from 2 to 10.5 with an interval of 0.5, three independent replicas under each pH condition were run for 50 ns. The Henderson–Hasselbalch equation was used to calculate the pKa values of titratable residues.

All subsequent simulations were performed using the GPU-accelerated version of GROMACS (v.2023) ^64^ with the CHARMM36m forcefield ^65^ with WYF parameters for cation-pi interactions ^66^, and the TIP3P water model. To study permeation through XPR1, Pi molecules were included within the system. As Pi exists in either the H_2_PO_4_^-^ or HPO_4_^2-^ states at pH 7, initial simulations were performed with 20 molecules of H_2_PO_4_^-^ and 20 molecules of HPO_4_^2-^ in each system. The structures of H_2_PO_4_^-^ and HPO_4_^2-^ were geometry optimised via quantum mechanics calculations using the Automated Topology Builder Webserver ^67^. Parameters for H_2_PO_4_^-^ and HPO_4_^2-^ were obtained using the CHARMM General Force Field (CGenFF) ^67^.

Steepest descent energy minimization was performed with a tolerance of 1,000 kJ mol^−1^ nm^−1^. During minimization, all protein backbone and sidechain atoms were position restrained with force-constants of 4,000 kJ mol^−1^ nm^−1^ and 2,000 kJ mol^−1^ nm^−1^, respectively. Position restraints of 1,000 kJ mol^−1^ nm^−1^ were also placed on the phosphorous atoms of DOPC headgroups. After minimizing, systems were equilibrated over six steps for a total of 60 ns where position restraints were gradually released on all atoms, with only the Cα atoms of residues 618-625 remaining restrained with a force-constant of 50 kJ mol^−1^ nm^−1^ to maintain the orientation of the C-Loop. The LINCS algorithm was used to constrain covalent bonds to hydrogen atoms, allowing a 2 fs timestep to be used ^68^.

Electrostatic interactions within a cut-off of 1.2 nm were calculated using the Particle-Mesh Ewald algorithm ^69^, using the Verlet grid cut-off scheme with an update frequency of every 20 steps. Long-range van der Waals interactions were calculated within a cut-off of 1.2 nm, with a smooth switching function beginning from 1.0 nm. All simulations were performed under periodic boundary conditions. In the equilibration stages, the Berendsen thermostat was used to maintain the simulation temperature at 310.15. K, using a time constant of 1 ps ^70^. Three thermally coupled groups consisting of 1) the protein, 2) the membrane, and 3) the water, Na^+^, Cl^-^, and Pi ions were used to improve the accuracy of the simulated temperature. The Berendsen barostat ^70^ was used to maintain simulation pressure at 1 bar, using a time constant of 5 ps with a compressibility of 4.5e-5 bar^-1^. During production runs, the v-rescale thermostat ^71^ and Parrinello-Rahman barostat ^72^ were used to maintain the temperature and pressure at 310.15 K and 1.0 bar, respectively.

To improve the chances of observing Pi permeation, an electric field was applied during production simulations along the Z-axis of the system with strengths of –0.035 and 0.035 V nm^−1^, to create a potential difference across the simulation box of –400 and 400 mV, respectively. These initial systems were simulated with five replicates for 500 ns each. Further simulations of both the iC-eC and iO-eO structures were performed using only H_2_PO_4_^-^, with 60 molecules per system. Five replicates of each system were simulated for 2 μs each at both –400 and 400 mV.

Measurements of the transport pathway radius were performed using an implementation of HOLE2.0 ^56^ within the MDAnalysis library ^73^. Analysis of Pi permeation events were performed using a script modified from ^74^. Measurement of Pi density along the Z-axis of the protein was analysed using GMX density. Systems were visualized using the Visual Molecular Dynamics software ^57^. The volumetric density of Pi within the cavity was obtained using the VMD VolMap plugin tool. All other analysis and graphing were performed in python using the MDAnalysis ^73^, Pandas ^75^, NumPy ^76^, Matplotlib ^77^, and Seaborn libraries ^78^. Statistical analysis was performed with SciPy ^79^.

## Statistics and Reproducibility

The cryo-EM micrographs were recorded with reproducible quality during data collection. The number of micrographs in each dataset is summarized in Extended Data Table 1, 2. Each protein purification was repeated at least three times with reproducible gel filtration profiles and SDS-PAGE band positions.

## Data availability

The following Cryo-EM maps and atomic coordinates have been deposited to EMDataBank and Protein Data Bank: XPR1-apo (EMD-xxx, PDB-xxx), XPR1-apo-SPX1 (EMD-xxx, PDB-xxx), XPR1-apo-SPX2 (EMD-xxx, PDB-xxx), XPR1-apo-SPX3 (EMD-xxx, PDB-xxx), XPR1-InsP8-SPX1 (EMD-xxx, PDB-xxx), XPR1-InsP8-SPX2 (EMD-xxx, PDB-xxx), XPR1-iC-eC (EMD-xxx, PDB-xxx), XPR1-iC-eO (EMD-xxx, PDB-xxx), XPR1-iO-eC (EMD-xxx, PDB-xxx), XPR1-iO-eO (EMD-xxx, PDB-xxx), XPR1-G621A (EMD-xxx, PDB-xxx), XPR1-R459C (EMD-xxx, PDB-xxx), XPR1-R570L (EMD-xxx, PDB-xxx), PXo (EMD-xxx), PXo-G604A (EMD-xxx, PDB-xxx), PHO1 (EMD-xxx), XPR1-KIDINS220 (EMD-xxx, PDB-xxx), XPR1-KIDINS220-InsP8 (EMD-xxx, PDB-xxx). Further data that support the findings of this study are available from the corresponding authors upon reasonable request.

## Author contributions

X.W. and Z.B. performed protein purification, cryo-EM sample preparation, data collection, image processing, and functional assay. X.W., Z.B., Y.S., and Y.Z. performed structural analysis. Y.H. assisted the data collection and structural analysis. C.W., R.J., and B.C. performed molecular dynamics simulations. H.J. sythineised InsP8. T.Y., M.L., H.W., C.G. and S.S. assisted in functional studies. Y.Z., B.C., and Y.S., and S.S.conceived and supervised the project. All authors wrote and approved the manuscript.

## Acknowledgement

We thank the Cryo-Electron Microscopy center at the Interdisciplinary Research Center on Biology and Chemistry, Shanghai Institute of Organic Chemistry for help with data collection. YZ is supported by STI2030-Major Projects (2022ZD0207400), Shanghai Key Laboratory of Aging Studies (19DZ2260400), and The Basic Research Pioneer Project by the Science and Technology Commission of Shanghai Municipality (STCSM). RJ and BC are supported by an Australian Research Council discovery project (DP200100860) This research/project was undertaken with the assistance of resources and services from the National Computational Infrastructure (NCI) and the Pawsey Supercomputing Research Centre, which are supported by the Australian Government and the Australian Government and the Government of Western Australia respectively. HW, CG, and SS are supported by the Intramural Research Program of the NIH, National Institute of Environmental Health Sciences (NIEHS).

## Competing interests

The authors declare no competing interests.

## Supplementary Figures

**Figure S1.**
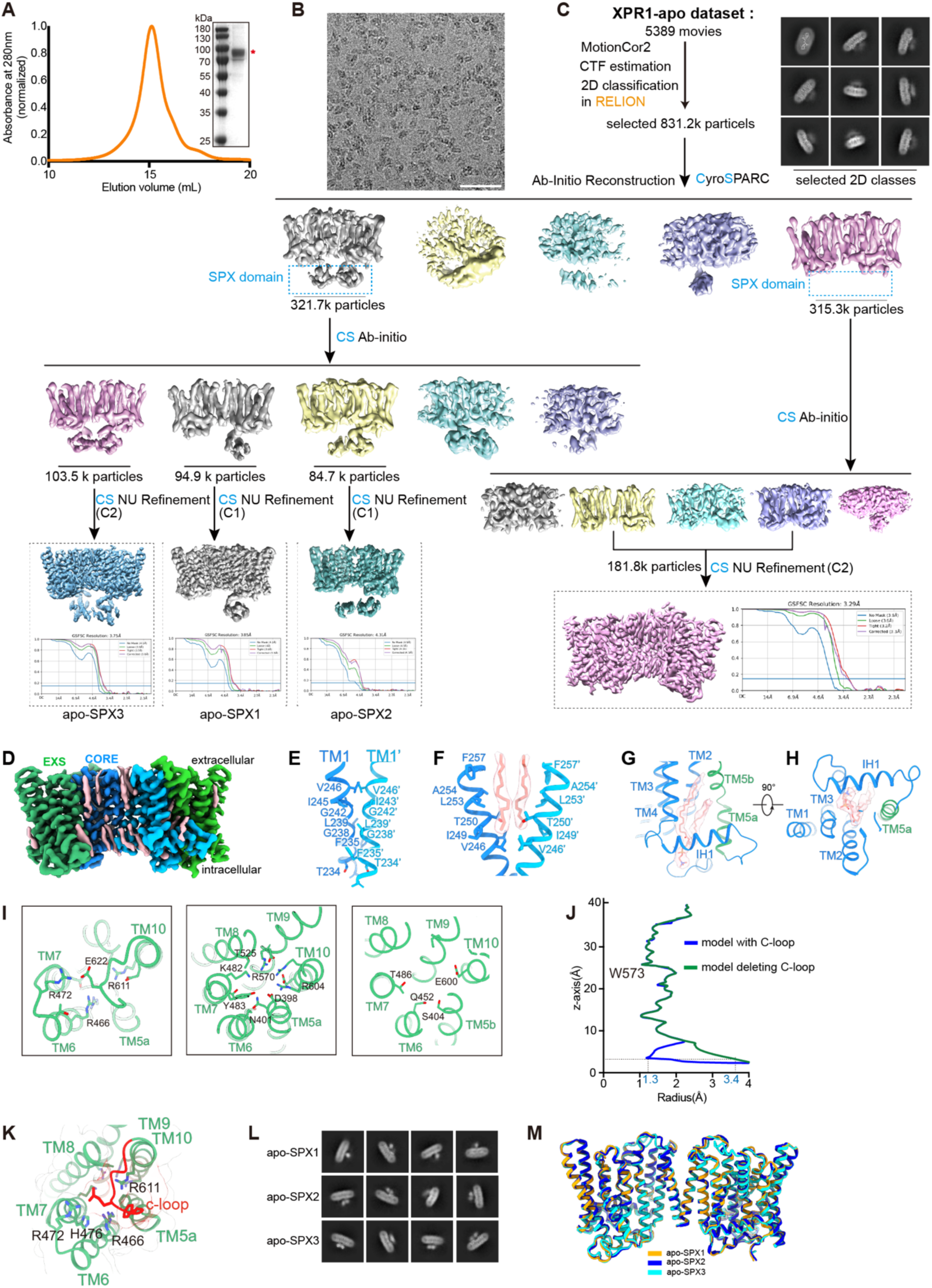
Protein purification and structure determination of XPR1 in apo state. **A**, Gel filtration profile of XPR1 with Coomassie blue staining of the peak fraction inset. Red sticks indicate the band of XPR1. **B**, An area of a representative cryo-EM micrograph of XPR1 in the apo state (scale bar: 50 nm). **C**, Image processing of XPR1 in the apo state with RELION and CryoSPARC. The map and FSC curve obtained by NU refinement are displayed in dashed boxes. **D**, The Cryo-EM map of XPR1 in the apo state is shown in the side view. **E**, TM1 forms the dimerization interface of XPR1. The interacting residues are shown as sticks, with the backbones of G238 and G242 involved in the interaction. **F**, The lipid acyl chain (transparent density with stick model) bridges TM1 dimerization, shown in side view. **G-H**, A zoomed-in view of the lipid (transparent density) sandwiched between IH1 and the transmembrane domain, shown in side (G) and bottom (H) views. **I**, The charged and polar residues in the center of the EXS domain are displayed as sticks. **J**, Radius plot of the permeation pathway of XPR1 modeled with (blue) and without (green) the C-loop residues. **K**, The C-loop covers the positively charged residues at the intracellular gate, with the surface of the C-loop highlighted in black silhouette. **L**, 2D averages of SPX in apo-SPX1, apo-SPX2, and apo-SPX3 states. **M**, Superimposing the models of XPR1 in apo-SPX1, apo-SPX2, and apo-SPX3 states indicates their similarity.

**Figure S2.**
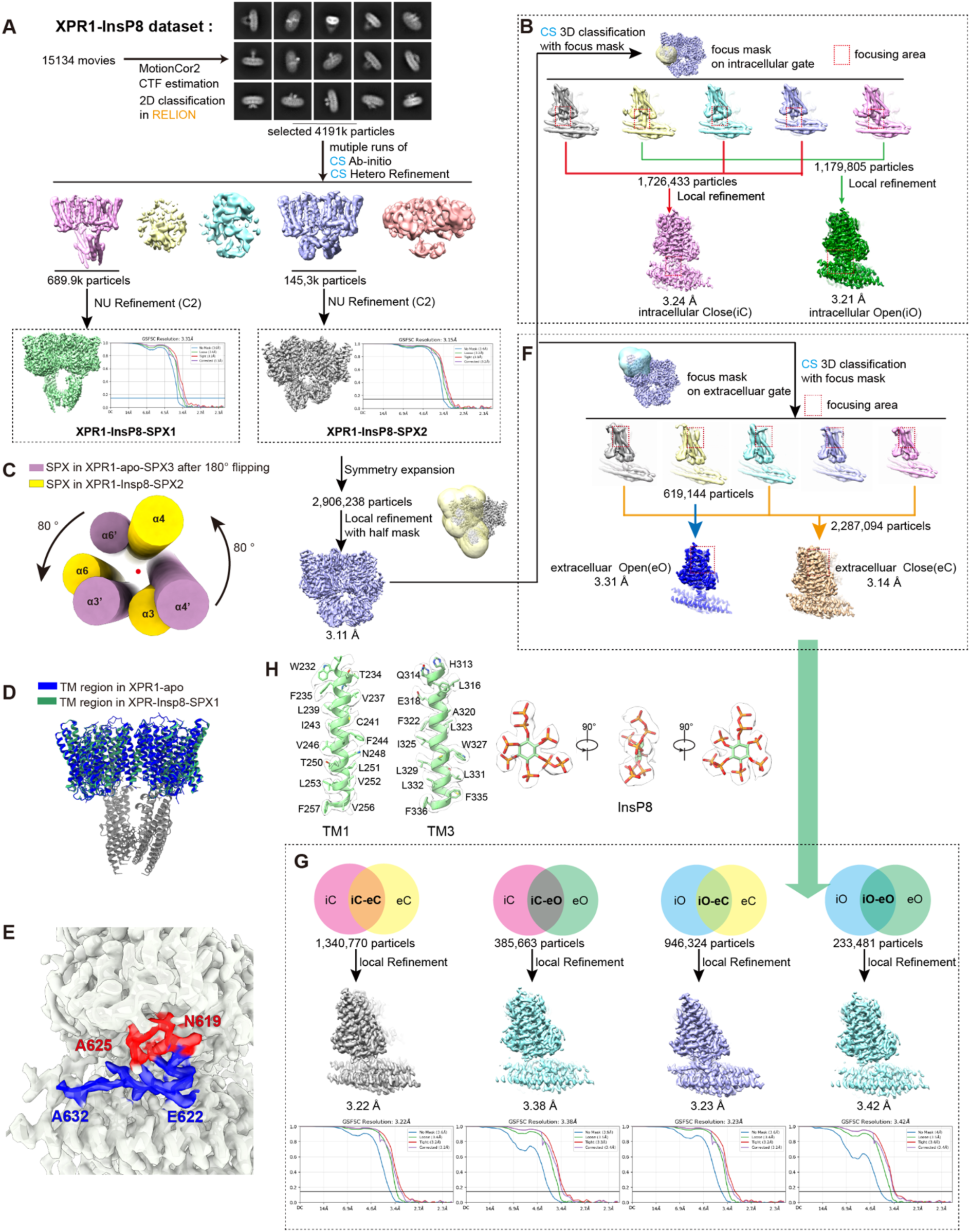
Data processing workflow of XPR1-InsP8 complex. **A**, Image processing of the XPR1-InsP8 complex with RELION and CryoSPARC. The map and FSC curve obtained by NU refinement are displayed in dashed boxes. **B**, Focused classification of XPR1-InsP8-SPX2 with a mask around the intracellular gate. **C**, The SPX domain rotates 80° along the elongated SPX axis (indicated by a red point) after 180° flipping along the axis perpendicular to the membrane. **D**, Superimposing the models of XPR1-apo and XPR1-InsP8-SPX2 states indicates their similarity. **E**, Cryo-EM map of XPR1-InsP8-SPX2 around the intracellular gate, showing a mixture of two different conformations (red and blue) of the C-loop. **F**, Focused classification of XPR1-InsP8-SPX2 with a mask around the extracellular gate. **G**, The strategy used to obtain the high-resolution structures of iC-eC, iO-eC, iC-eO, and iO-eO states with overlapping particles. Gold standard FSC curves (FSC = 0.143) are displayed below the refined maps. **H**, The local density of TM1, TM3, and InsP8 in the N-C pocket was shown to indicate map quality.

**Figure S3.**
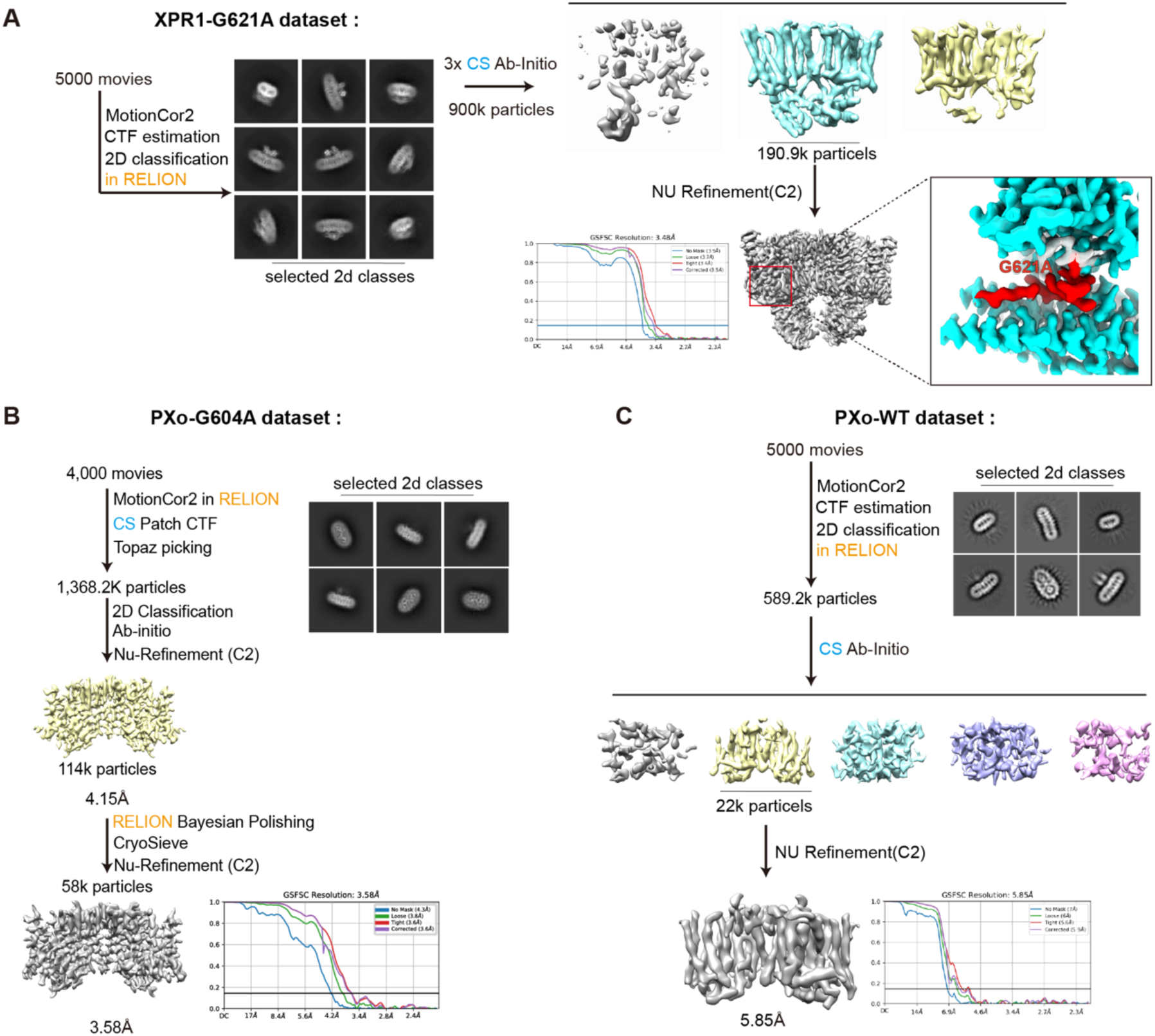
Data processing of XPR1-G621A, PXo-WT, and PXo-G604A. **A**, Image processing of XPR1-G621A with RELION and CryoSPARC. A cryo-EM map of XPR1-G621A around the intracellular gate, showing the release of the C-loop (red), is displayed in a zoomed-in view. **B-C**, Image processing of PXo-WT (C) and PXo-G604A (B) with RELION and CryoSPARC.

**Figure S4.**
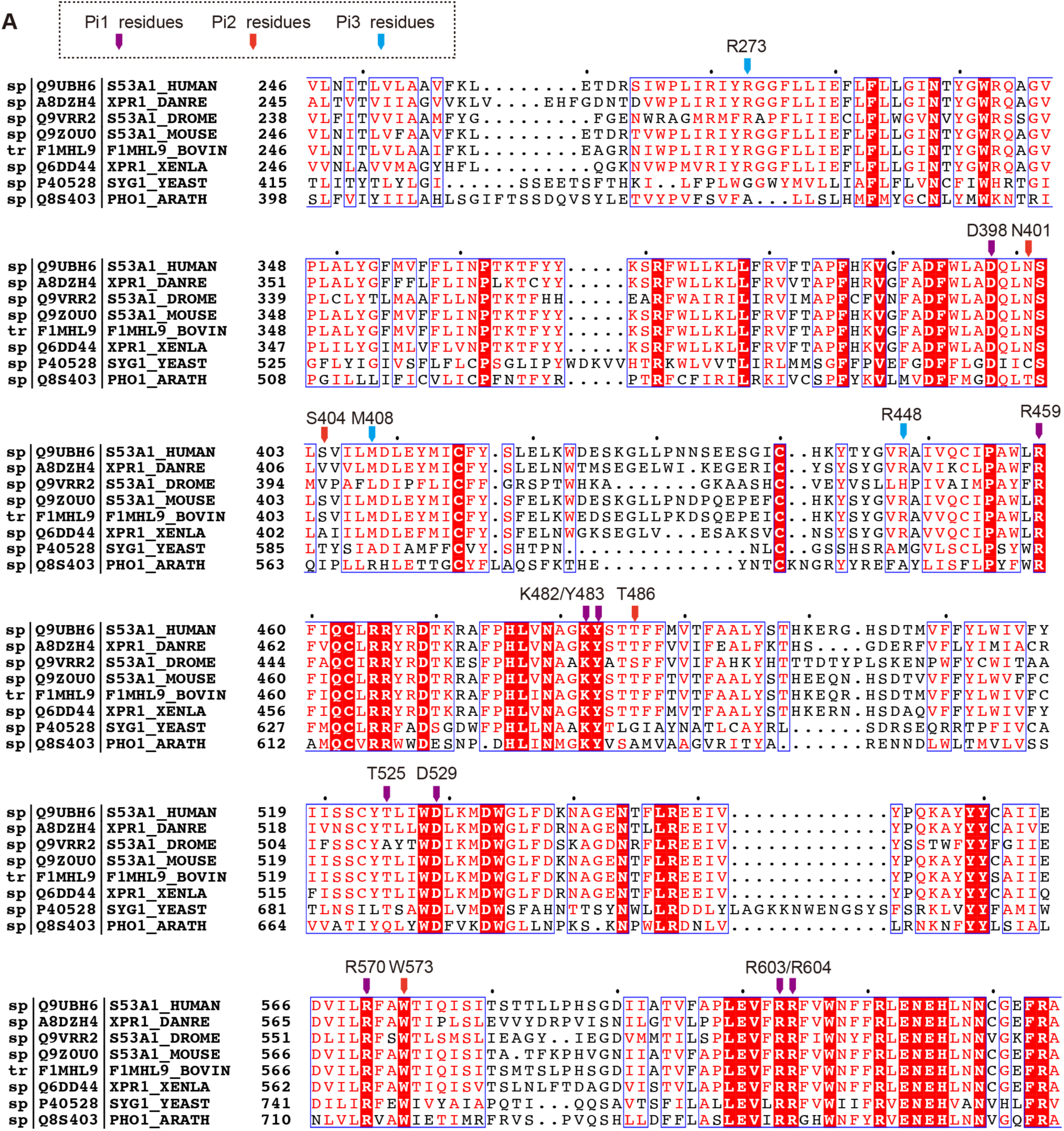
Sequence alignment of XPR1 orthologues. **A**, Sequence alignment of XPR1 and orthologues. The pore residues involved in Pi recognition are indicated by purple arrows (for Pi-1 residues), orange arrows (for Pi-2 residues), and blue arrows (for Pi-3 residues).

**Figure S5.**
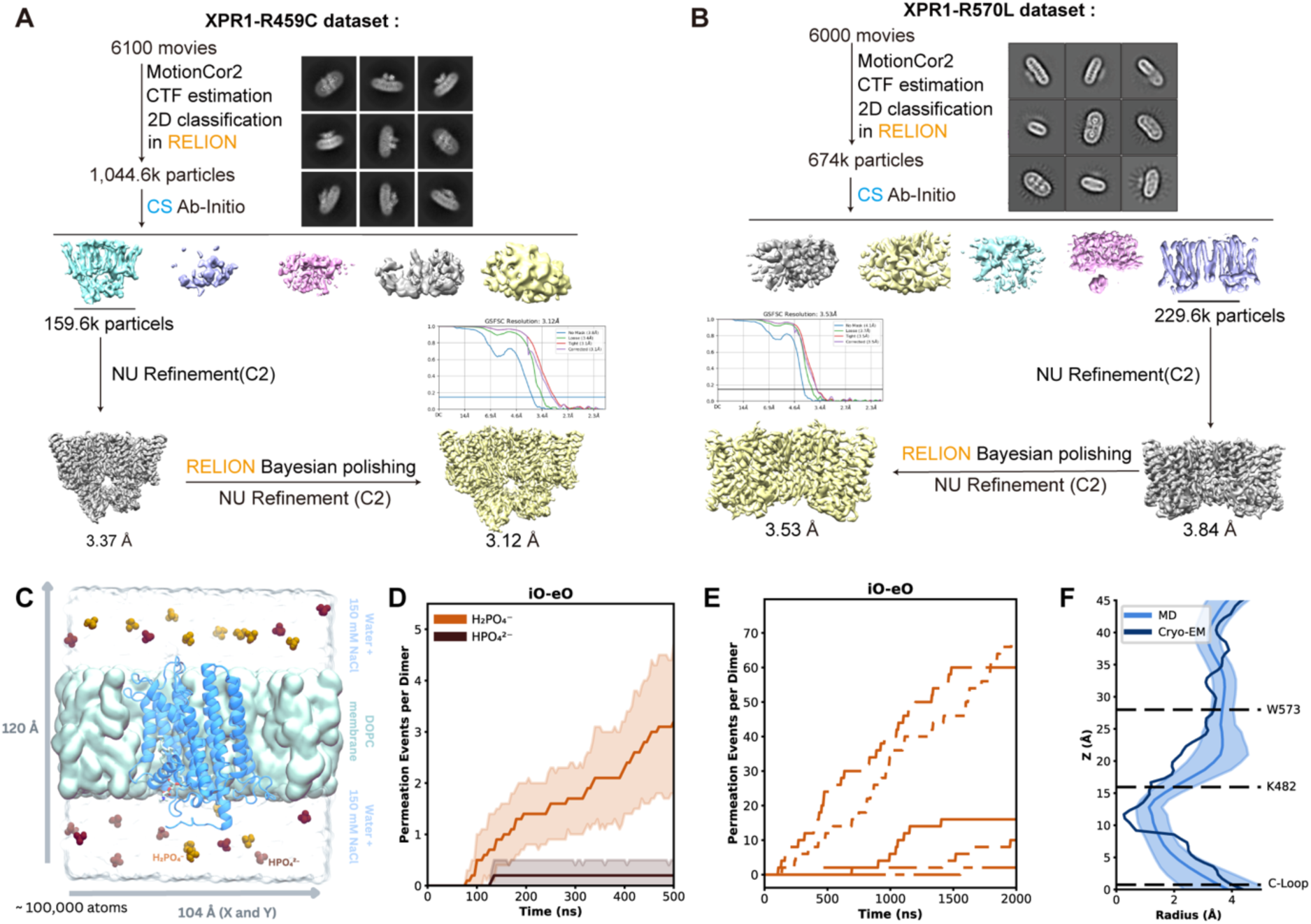
Data processing of XPR1 disease mutants and MD simulation of XPR1 in iO-eO state. **A-B**, Image processing of XPR1-R459C (A) and XPR1-R570L (B) with RELION and CryoSPARC. See methods and Table S1 for details. **C**, Simulation system containing XPR1 iO-eO monomer (blue) embedded in a DOPC bilayer (teal surface). The system is solvated in water (transparent surface), 150mM NaCl (not shown) and in this example 12 H₂PO₄⁻ (light orange) and 12HPO₄²⁻ (dark orange) molecules. **D**, Cumulative H₂PO₄⁻ (light orange) and HPO₄²⁻ (dark brown) permeation events recorded from four replicates of MD simulations of the iO-eO structure at –400 mV. **E**, Cumulative Pi permeation events recorded from four replicates of MD simulations of the iO-eO structure containing just the H₂PO₄⁻ species of phosphate at –400 mV. **F**, Comparison of the pore radius in the iO-eO cryo-EM structure to the average pore radius (± SEM) measured from simulations. The C-Loop, K482, and W573 positions are given as a reference.

**Figure S6.**
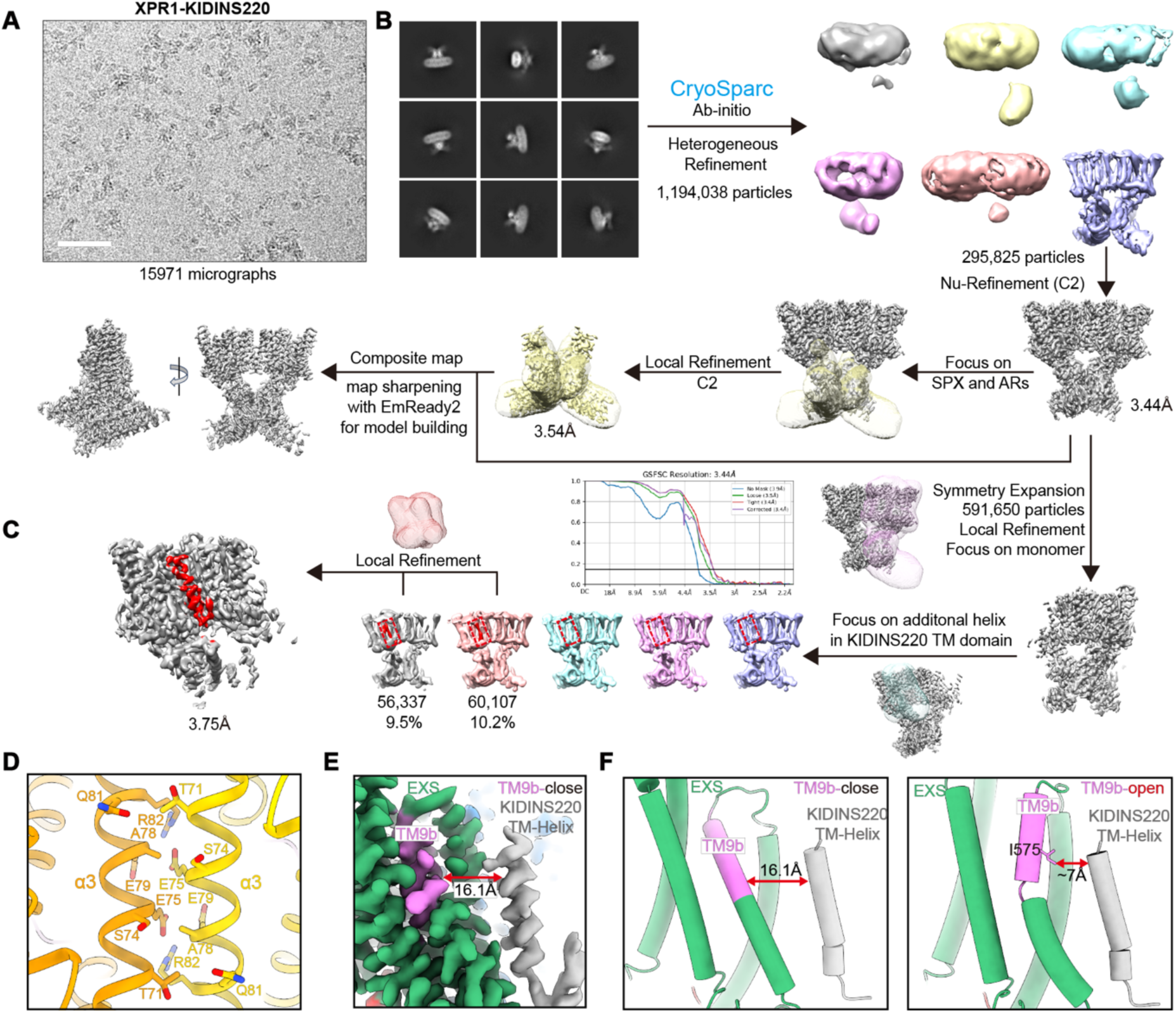
Data processing workflow and structure of the XPR1-KIDINS220 complex. **A**, An area of a representative cryo-EM micrograph of XPR1-KIDINS220 (scale bar: 50 nm). **B**, Image processing of XPR1-KIDINS220 with RELION and CryoSPARC. **C**, Focused classification of XPR1-KIDINS220 with a mask around the additional TM helix. **D**, SPX α3 helix dimerization interface in the XPR1-KIDINS220 complex. The interacting residues are shown as sticks. **E**, The location of the KIDINS220 TM helix (grey) on the EXS domain (green). TM9b is colored violet. **F**, The distance between the KIDINS220 TM helix (grey) and the TM9b helix (violet) in the extracellular gate closed (left) and open (right) states. The distance is measured between the Cα of the tip residue on the KIDINS220 TM helix and the side chain of ILE575 on TM9b.

**Figure S7.**
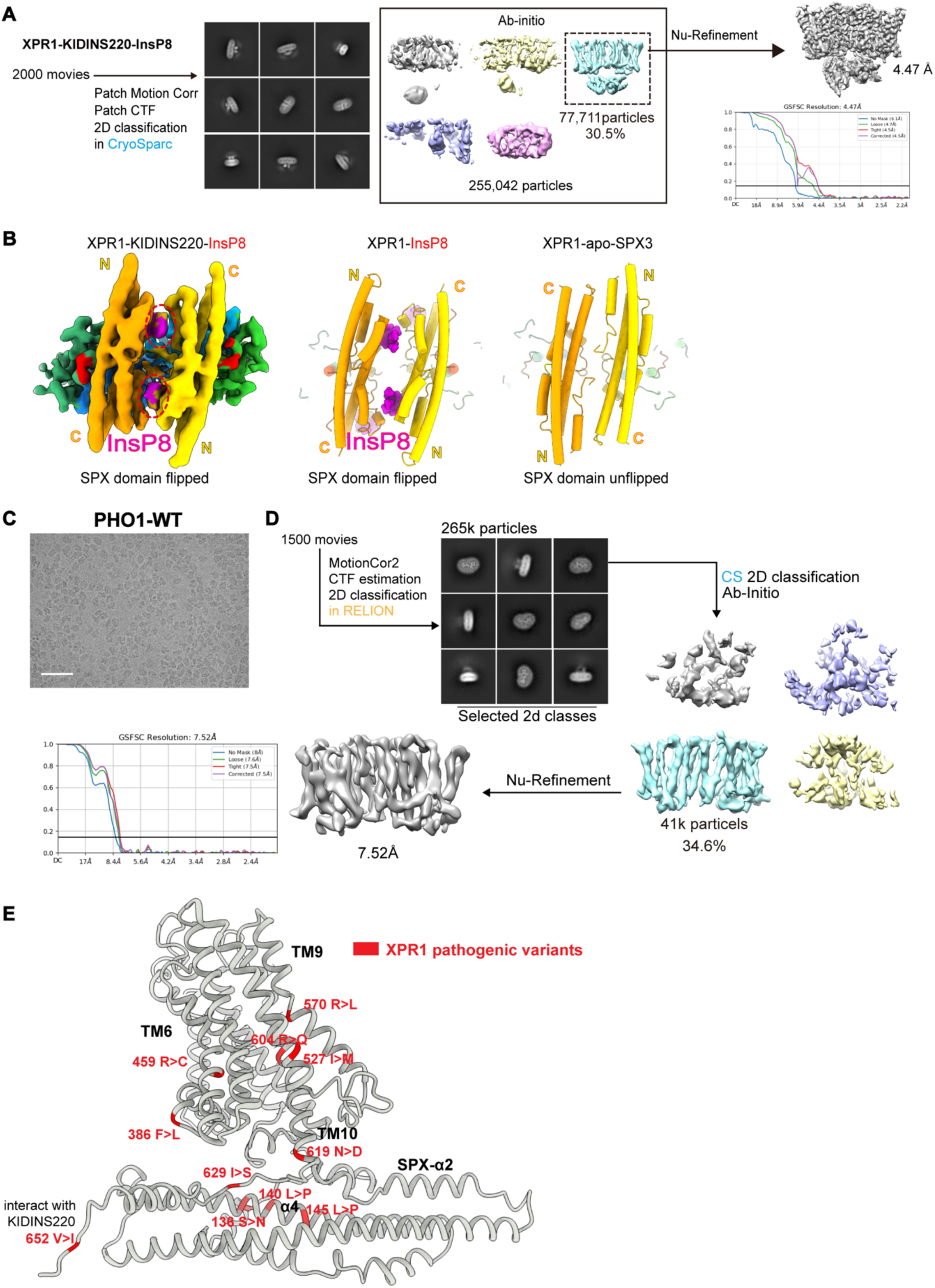
Data processing workflow and structure of the XPR1-KIDINS220-InsP8 complex and PHO1. **A**, Image processing of the XPR1-KIDINS220-InsP8 complex. The gold standard FSC curve (FSC = 0.143) calculated with NU refinement is displayed with the refined map. **B**, The orientation of the SPX domain in the XPR1-KIDINS220-InsP8 complex (left) is the same as in the XPR1-InsP8 complex (middle) but different from that in XPR1-apo-SPX3 (right). **C**, An area of a representative cryo-EM micrograph of PHO1 (scale bar: 50 nm). **D**, Image processing of PHO1 with RELION and CryoSPARC. See methods and Table S1 for details. **E**, Mapping of PFBC pathogenic variants on XPR1 subunit (grey), with variants labeled in red. R459, R570, and R604 are located in the permeation pathway. N619 and I629 are located at the intracellular gate. V652 is located at the flexible distal C-terminal end, but it is involved in KIDINS220 interaction (Fig. 5E).

**Table S1.** Cryo-EM Data collection, refinement and validation statistics-part1. This table provides comprehensive information on the Cryo-EM data collection, refinement, and validation statistics for the studied samples.

|  | XPR1-apo | XPR1-apo-SPX1 | XPR1-apo-SPX2 | XPR1-apo-SPX3 | XPR1-InsP8-SPX1 | XPR1-InsP8-SPX2 | XPR1-iC-eC | XPR1-iC-eO | XPR1-iO-eC | XPR1-iO-eO |
| --- | --- | --- | --- | --- | --- | --- | --- | --- | --- | --- |
|  | (EMDB-)<br>(PDB xxx) | (EMDB-)<br>(PDB xxx) | (EMDB-)<br>(PDB xxx) | (EMDB-)<br>(PDB xxx) | (EMDB-)<br>(PDB xxx) | (EMDB-)<br>(PDB xxx) | (EMDB-)<br>(PDB xxx) | (EMDB-)<br>(PDB xxx) | (EMDB-)<br>(PDB xxx) | (EMDB-)<br>(PDB xxx) |
| Data collection and processing |  |  |  |  |  |  |  |  |  |  |
| Magnification | 81,000 | 81,000 | 81,000 | 81,000 | 81,000 | 81,000 | 81,000 | 81,000 | 81,000 | 81,000 |
| Voltage (kV) | 300 | 300 | 300 | 300 | 300 | 300 | 300 | 300 | 300 | 300 |
| Electron exposure (e<sup>-</sup>/Å<sup>2</sup>) | 49.41 | 49.41 | 49.41 | 49.41 | 49.41 | 49.41 | 49.41 | 49.41 | 49.41 | 49.41 |
| Defocus range (µm) | -1.4 ~ -2.4 | -1.4 ~ -2.4 | -1.4 ~ -2.4 | -1.4 ~ -2.4 | -1.4 ~ -2.4 | -1.4 ~ -2.4 | -1.4 ~ -2.4 | -1.2 ~ -2.4 | -1.4 ~ -2.4 | -1.4 ~ -2.4 |
| Pixel size (Å) | 1.055 | 1.055 | 1.055 | 1.055 | 1.055 | 1.055 | 1.055 | 1.055 | 1.055 | 1.055 |
| Symmetry imposed | C2 | C1 | C1 | C2 | C2 | C2 | C1 | C1 | C1 | C1 |
| Initial particle images (no.) | 831191 | 831191 | 831191 | 831191 | 4191317 | 4191317 | 2906238 | 2906238 | 2906238 | 2906238 |
| Final particle images (no.) | 181792 | 94997 | 84735 | 103528 | 694096 | 1452119 | 1340770 | 385663 | 946324 | 233481 |
| Map resolution (Å) | 3.29 | 3.85 | 4.31 | 3.75 | 3.31 | 3.15 | 3.22 | 3.38 | 3.23 | 3.48 |
| FSC threshold | 0.143 | 0.143 | 0.143 | 0.143 | 0.143 | 0.143 | 0.143 | 0.143 | 0.143 | 0.143 |
| Map resolution range (Å) | 3.0-5.0 | 3.5-10.0 | 4.0-10.0 | 3.0-15 | 3.3-5.0 | 3.0-3.5 | 3.1-3.5 | 3.2-3.5 | 3.1-3.5 | 3.1-3.5 |
| Refinement |  |  |  |  |  |  |  |  |  |  |
| Initial model used (PDB code) | AF-Q9UBH6-F1 | AF-Q9UBH6-F1 | AF-Q9UBH6-F1 | AF-Q9UBH6-F1 | AF-Q9UBH6-F1 | AF-Q9UBH6-F1 | AF-Q9UBH6-F1 | AF-Q9UBH6-F1 | AF-Q9UBH6-F1 | AF-Q9UBH6-F1 |
| Map sharpening *B* factor (Å<sup>2</sup>) | 146.8 | 127.7 | 191.5 | 145.0 | 180.9 | 201.3 | 181.0 | 153.0 | 174.9 | 143.5 |
| Model composition |  |  |  |  |  |  |  |  |  |  |
| Non-hydrogen atoms | 6304 | 7588 | 9181 | 9676 | 9338 | 9976 | 4961 | 5011 | 5018 | 5063 |
| Protein residues | 754 | 906 | 1095 | 1160 | 1094 | 1178 | 596 | 602 | 604 | 609 |
| Ligands | 0 | 0 | 0 | 0 | 4 | 4 | 3 | 2 | 3 | 2 |
| R.m.s. deviations |  |  |  |  |  |  |  |  |  |  |
| Bond lengths (Å) | 0.006 | 0.005 | 0.004 | 0.013 | 0.012 | 0.012 | 0.007 | 0.006 | 0.006 | 0.008 |
| Bond angles (°) | 0.757 | 0.636 | 0.638 | 1.066 | 1.024 | 1.470 | 0.878 | 0.942 | 0.747 | 0.721 |
| Validation |  |  |  |  |  |  |  |  |  |  |
| MolProbity score | 2.17 | 1.94 | 1.84 | 2.16 | 2.30 | 2.41 | 2.13 | 2.20 | 2.06 | 2.02 |
| Clash score | 14.61 | 10.44 | 9.44 | 11.59 | 19.62 | 21.60 | 14.37 | 19.42 | 12.29 | 10.59 |
| Poor rotamers (%) | 0 | 0.50 | 0 | 0.58 | 0.41 | 0.77 | 0.19 | 0.56 | 0.19 | 0 |
| Ramachandran plot |  |  |  |  |  |  |  |  |  |  |
| Outliers (%) | 0 | 0 | 0.09 | 0.96 | 0.19 | 0.69 | 0.19 | 0.50 | 0 | 0.17 |
| Allowed (%) | 8.36 | 5.94 | 4.85 | 10.14 | 8.60 | 10.52 | 7.46 | 5.85 | 7.36 | 7.44 |
| Favored (%) | 91.64 | 94.04 | 95.06 | 88.90 | 91.21 | 88.79 | 92.54 | 93.65 | 92.64 | 92.40 |

|  | XPR1-G621A | XPR1-R459C | XPR1-R570L | PXo-G604A | XPR1-KIDINS220 |
| --- | --- | --- | --- | --- | --- |
|  | (EMDB-)<br>(PDB xxx) | (EMDB-)<br>(PDB xxx) | (EMDB-)<br>(PDB xxx) | (EMDB-)<br>(PDB xxx) | (EMDB-)<br>(PDB xxx) |
| Data collection and processing |  |  |  |  |  |
| Magnification | 81,000 | 81,000 | 81,000 | 81,000 | 81,000 |
| Voltage (kV) | 300 | 300 | 300 | 300 | 300 |
| Electron exposure (e<sup>-</sup>/Å<sup>2</sup>) | 49.41 | 49.41 | 49.41 | 49.41 | 49.41 |
| Defocus range (µm) | -1.4 ~ -2.4 | -1.4 ~ -2.4 | -1.4 ~ -2.4 | -1.4 ~ -2.4 | -1.4 ~ -2.4 |
| Pixel size (Å) | 1.055 | 1.055 | 1.055 | 1.055 | 1.055 |
| Symmetry imposed | C2 | C2 | C2 | C2 | C2 |
| Initial particle images (no.) | 900042 | 1044666 | 674459 | 1,368,175 | 3,458,119 |
| Final particle images (no.) | 190974 | 159655 | 229444 | 58,767 | 295,825 |
| Map resolution (Å) | 3.48 | 3.12 | 3.53 | 3.53 | 3.44 |
| FSC threshold | 0.143 | 0.143 | 0.143 | 0.143 | 0.143 |
| Map resolution range (Å) | 3.2-3.5 | 3.0-3.5 | 3.2-3.8 | 2.8-5.0 | 2.9-4.7 |
| Refinement |  |  |  |  |  |
| Initial model used (PDB code) | AF-Q9UBH6-F1 | AF-Q9UBH6-F1 | AF-Q9UBH6-F1 | AF-Q9VRR2-F1 | AF-Q9ULH0-F1 |
| Map sharpening *B* factor (Å<sup>2</sup>) | 152.1 | 123.6 | 172.3 | -134.7 | -124.4 |
| Model composition |  |  |  |  |  |
| Non-hydrogen atoms | 9906 | 9748 | 6430 | 11290 | 16094 |
| Protein residues | 1192 | 1174 | 770 | 1410 | 1998 |
| Ligands | 0 | 0 | 0 | 0 | 0 |
| R.m.s. deviations |  |  |  |  |  |
| Bond lengths (Å) | 0.010 | 0.005 | 0.005 | 0.041 | 0.006 |
| Bond angles (°) | 0.904 | 0.777 | 0.704 | 3.375 | 1.263 |
| Validation |  |  |  |  |  |
| MolProbity score | 2.18 | 1.82 | 2.08 | 1.08 | 2.04 |
| Clash score | 14.15 | 11.40 | 11.22 | 1.36 | 12.41 |
| Poor rotamers (%) | 0 | 0 | 0 | 0.60 | 0.17 |
| Ramachandran plot |  |  |  |  |  |
| Outliers (%) | 0.38 | 0 | 0 | 0 | 0.46 |
| Allowed (%) | 8.93 | 3.70 | 8.66 | 3.40 | 6.27 |
| Favored (%) | 91.07 | 96.30 | 91.34 | 96.60 | 93.28 |

**Video S1: The structure of XPR1-KIDINS220 complex and conformational landscape of XPR1 activation**.

The video shows the structure of XPR1-KIDINS220 complex and the model of XPR1 activation.

## Notes

### Competing Interest Statement

The authors have declared no competing interest.

## References

1. Berndt, T., and Kumar, R. (2007). Phosphatonins and the regulation of phosphate homeostasis. Annu. Rev. Physiol. 69. 10.1146/annurev.physiol.69.040705.141729.

2. Peacock, M. (2021). Phosphate metabolism in health and disease. Calcified tissue international 108, 3–15.

3. Wagner, C.A. (2024). The basics of phosphate metabolism. Nephrology Dialysis Transplantation 39, 190–201.

4. Razzaque, M.S. (2011). Phosphate toxicity: new insights into an old problem. Clinical science 120, 91–97.

5. Forster, I.C., Hernando, N., Biber, J., and Murer, H. (2013). Phosphate transporters of the SLC20 and SLC34 families. Molecular aspects of medicine 34, 386–395.

6. Levi, M., Gratton, E., Forster, I.C., Hernando, N., Wagner, C.A., Biber, J., Sorribas, V., and Murer, H. (2019). Mechanisms of phosphate transport. Nature Reviews Nephrology 15, 482–500.

7. Tsai, J.-Y., Chu, C.-H., Lin, M.-G., Chou, Y.-H., Hong, R.-Y., Yen, C.-Y., Hsiao, C.-D., and Sun, Y.-J. (2020). Structure of the sodium-dependent phosphate transporter reveals insights into human solute carrier SLC20. Science Advances 6, eabb4024.

8. Giovannini, D., Touhami, J., Charnet, P., Sitbon, M., and Battini, J.-L. (2013). Inorganic phosphate export by the retrovirus receptor XPR1 in metazoans. Cell Rep 3, 1866–1873.

9. Wege, S., and Poirier, Y. (2014). Expression of the mammalian xenotropic polytropic virus receptor 1 (XPR1) in tobacco leaves leads to phosphate export. FEBS Lett. 588. 10.1016/j.febslet.2013.12.013.

10. Jennings, M.L. (2023). Role of transporters in regulating mammalian intracellular inorganic phosphate. Front. Pharmacol. 14. 10.3389/fphar.2023.1163442.

11. Battini, J.L., Rasko, J.E., and Miller, A.D. (1999). A human cell-surface receptor for xenotropic and polytropic murine leukemia viruses: possible role in G protein-coupled signal transduction. Proc. Natl Acad. Sci. USA 96. 10.1073/pnas.96.4.1385.

12. Tailor, C.S., Nouri, A., Lee, C.G., Kozak, C., and Kabat, D. (1999). Cloning and characterization of a cell surface receptor for xenotropic and polytropic murine leukemia viruses. Proceedings of the National Academy of Sciences 96, 927–932.

13. Yang, Y.L. (1999). Receptors for polytropic and xenotropic mouse leukaemia viruses encoded by a single gene at Rmc1. Nat. Genet. 21. 10.1038/6005.

14. Li, X., Gu, C., Hostachy, S., Sahu, S., Wittwer, C., Jessen, H.J., Fiedler, D., Wang, H., and Shears, S.B. (2020). Control of XPR1-dependent cellular phosphate efflux by InsP8 is an exemplar for functionally-exclusive inositol pyrophosphate signaling. Proceedings of the National Academy of Sciences 117, 3568–3574.

15. Moritoh, Y., Abe, S.-i., Akiyama, H., Kobayashi, A., Koyama, R., Hara, R., Kasai, S., and Watanabe, M. (2021). The enzymatic activity of inositol hexakisphosphate kinase controls circulating phosphate in mammals. Nature communications 12, 4847.

16. Gadsby, D.C., Vergani, P., and Csanady, L. (2006). The ABC protein turned chloride channel whose failure causes cystic fibrosis. Nature 440. 10.1038/nature04712.

17. Legati, A., Giovannini, D., Nicolas, G., López-Sánchez, U., Quintáns, B., Oliveira, J.R., Sears, R.L., Ramos, E.M., Spiteri, E., and Sobrido, M.-J. (2015). Mutations in XPR1 cause primary familial brain calcification associated with altered phosphate export. Nature genetics 47, 579–581.

18. Anheim, M. (2016). XPR1 mutations are a rare cause of primary familial brain calcification. J. Neurol. 263. 10.1007/s00415-016-8166-4.

19. Ansermet, C. (2017). Renal Fanconi syndrome and hypophosphatemic rickets in the absence of xenotropic and polytropic retroviral receptor in the nephron. J. Am. Soc. Nephrol. 28. 10.1681/ASN.2016070726.

20. Yao, X.-P., Zhao, M., Wang, C., Guo, X.-X., Su, H.-Z., Dong, E.-L., Chen, H.-T., Lai, J.-H., Liu, Y.-B., and Wang, N. (2017). Analysis of gene expression and functional characterization of XPR1: a pathogenic gene for primary familial brain calcification. Cell and tissue research 370, 267–273.

21. López-Sánchez, U., Nicolas, G., Richard, A.-C., Maltête, D., Charif, M., Ayrignac, X., Goizet, C., Touhami, J., Labesse, G., and Battini, J.-L. (2019). Characterization of XPR1/SLC53A1 variants located outside of the SPX domain in patients with primary familial brain calcification. Scientific reports 9, 6776.

22. Akasu-Nagayoshi, Y., Hayashi, T., Kawabata, A., Shimizu, N., Yamada, A., Yokota, N., Nakato, R., Shirahige, K., Okamoto, A., and Akiyama, T. (2022). PHOSPHATE exporter XPR1/SLC53A1 is required for the tumorigenicity of epithelial ovarian cancer. Cancer Science 113, 2034–2043.

23. Bondeson, D.P., Paolella, B.R., Asfaw, A., Rothberg, M.V., Skipper, T.A., Langan, C., Mesa, G., Gonzalez, A., Surface, L.E., and Ito, K. (2022). Phosphate dysregulation via the XPR1–KIDINS220 protein complex is a therapeutic vulnerability in ovarian cancer. Nature cancer 3, 681–695.

24. Orimo, K., Kakumoto, T., Hara, R., Goto, R., Ishiura, H., Mitsui, J., Yoshida, C., Uesaka, Y., Suzuki, Y., and Morishita, S. (2023). A Japanese family with idiopathic basal ganglia calcification carrying a novel XPR1 variant. Journal of the neurological sciences 451.

25. Hamburger, D., Rezzonico, E., MacDonald-Comber Petetot, J., Somerville, C., and Poirier, Y. (2002). Identification and characterization of the Arabidopsis PHO1 gene involved in phosphate loading to the xylem. Plant Cell 14. 10.1105/tpc.000745.

26. Secco, D., Wang, C., Shou, H., and Whelan, J. (2012). Phosphate homeostasis in the yeast Saccharomyces cerevisiae, the key role of the SPX domain-containing proteins. FEBS Lett. 586. 10.1016/j.febslet.2012.01.036.

27. Xu, C., Xu, J., Tang, H.-W., Ericsson, M., Weng, J.-H., DiRusso, J., Hu, Y., Ma, W., Asara, J.M., and Perrimon, N. (2023). A phosphate-sensing organelle regulates phosphate and tissue homeostasis. Nature 617, 798–806.

28. Wild, R. (2016). Control of eukaryotic phosphate homeostasis by inositol polyphosphate sensor domains. Science 352. 10.1126/science.aad9858.

29. Wilson, M.S., Livermore, T.M., and Saiardi, A. (2013). Inositol pyrophosphates: between signalling and metabolism. Biochemical Journal 452, 369–379.

30. Wege, S. (2016). The EXS domain of PHO1 participates in the response of shoots to phosphate deficiency via a root-to-shoot signal. Plant Physiol. 170. 10.1104/pp.15.00975.

31. López-Sánchez, U., Tury, S., Nicolas, G., Wilson, M.S., Jurici, S., Ayrignac, X., Courgnaud, V., Saiardi, A., Sitbon, M., and Battini, J.-L. (2020). Interplay between primary familial brain calcification-associated SLC20A2 and XPR1 phosphate transporters requires inositol polyphosphates for control of cellular phosphate homeostasis. Journal of Biological Chemistry 295, 9366–9378.

32. Li, X., Kirkpatrick, R.B., Wang, X., Tucker, C.J., Shukla, A., Jessen, H.J., Wang, H., Shears, S.B., and Gu, C. (2024). Homeostatic coordination of cellular phosphate uptake and efflux requires an organelle-based receptor for the inositol pyrophosphate IP8. Cell Rep 43.

33. Iglesias, T., Cabrera-Poch, N., Mitchell, M.P., Naven, T.J., Rozengurt, E., and Schiavo, G. (2000). Identification and cloning of Kidins220, a novel neuronal substrate of protein kinase D. Journal of Biological Chemistry 275, 40048–40056.

34. Cesca, F., Yabe, A., Spencer-Dene, B., Arrigoni, A., Al-Qatari, M., Henderson, D., Phillips, H., Koltzenburg, M., Benfenati, F., and Schiavo, G. (2011). Kidins220/ARMS is an essential modulator of cardiovascular and nervous system development. Cell death & disease 2, e226–e226.

35. Neubrand, V.E., Cesca, F., Benfenati, F., and Schiavo, G. (2012). Kidins220/ARMS as a functional mediator of multiple receptor signalling pathways. Journal of cell science 125, 1845–1854.

36. Capolicchio, S., Wang, H., Thakor, D.T., Shears, S.B., and Jessen, H.J. (2014). Synthesis of Densely Phosphorylated Bis-1, 5-Diphospho-myo-Inositol Tetrakisphosphate and its Enantiomer by Bidirectional P-Anhydride Formation. Angewandte Chemie 126, 9662–9665.

37. Yan, R., Chen, H., Liu, C., Zhao, J., Wu, D., Jiang, J., Gong, J., and Jiang, D. (2024). Human XPR1 structures reveal phosphate export mechanism. Nature, 1-8.

38. Scholz-Starke, J., and Cesca, F. (2016). Stepping out of the shade: control of neuronal activity by the scaffold protein Kidins220/ARMS. Frontiers in cellular neuroscience 10, 68.

39. Drew, D., and Boudker, O. (2016). Shared molecular mechanisms of membrane transporters. Annu. Rev. Biochem. 85. 10.1146/annurev-biochem-060815-014520.

40. Cheng, X., Zhao, M., Chen, L., Huang, C., Xu, Q., Shao, J., Wang, H.-T., Zhang, Y., Li, X., and Xu, X. (2024). Astrocytes modulate brain phosphate homeostasis via polarized distribution of phosphate uptake transporter PiT2 and exporter XPR1. Neuron.

41. Gamir-Morralla, A., Belbin, O., Fortea, J., Alcolea, D., Ferrer, I., Lleó, A., and Iglesias, T. (2017). Kidins220 correlates with Tau in alzheimer’s disease brain and cerebrospinal fluid. Journal of Alzheimer’s Disease 55, 1327–1333.

42. Lu, Y., Yue, C.-X., Zhang, L., Yao, D., Xia, Y., Zhang, Q., Zhang, X., Li, S., Shen, Y., Cao, M., et al. Structural basis for inositol pyrophosphate gating of the phosphate channel XPR1. Science 0, eadp3252. doi:10.1126/science.adp3252.

43. He, Q., Zhang, R., Tury, S., Courgnaud, V., Liu, F., Battini, J.-l., Li, B., and Chen, Q. (2024). Structural basis of phosphate export by human XPR1. bioRxiv, 2024.2008.2022.609128.

44. Liu, Z. (2024). Structural insights into the mechanism of phosphate recognition and transport by human XPR1. bioRxiv, 2024.2008.2019.608714.

45. Goehring, A., Lee, C.H., Wang, K.H., Michel, J.C., Claxton, D.P., Baconguis, I., Althoff, T., Fischer, S., Garcia, K.C., and Gouaux, E. (2014). Screening and large-scale expression of membrane proteins in mammalian cells for structural studies. Nat Protoc 9, 2574–2585. 10.1038/nprot.2014.173.

46. Zheng, S.Q., Palovcak, E., Armache, J.P., Verba, K.A., Cheng, Y., and Agard, D.A. (2017). MotionCor2: anisotropic correction of beam-induced motion for improved cryo-electron microscopy. Nat Methods 14, 331–332. 10.1038/nmeth.4193.

47. Rohou, A., and Grigorieff, N. (2015). CTFFIND4: Fast and accurate defocus estimation from electron micrographs. J Struct Biol 192, 216–221. 10.1016/j.jsb.2015.08.008.

48. Zivanov, J., Nakane, T., Forsberg, B.O., Kimanius, D., Hagen, W.J., Lindahl, E., and Scheres, S.H. (2018). New tools for automated high-resolution cryo-EM structure determination in RELION-3. Elife 7. 10.7554/eLife.42166.

49. Punjani, A., Rubinstein, J.L., Fleet, D.J., and Brubaker, M.A. (2017). cryoSPARC: algorithms for rapid unsupervised cryo-EM structure determination. Nat Methods 14, 290–296. 10.1038/nmeth.4169.

50. Zhu, J., Zhang, Q., Zhang, H., Shi, Z., Hu, M., and Bao, C. (2023). A minority of final stacks yields superior amplitude in single-particle cryo-EM. Nature Communications 14, 7822. 10.1038/s41467-023-43555-x.

51. Bepler, T., Morin, A., Rapp, M., Brasch, J., Shapiro, L., Noble, A.J., and Berger, B. (2019). Positive-unlabeled convolutional neural networks for particle picking in cryo-electron micrographs. Nature methods 16, 1153–1160.

52. He, J., Li, T., and Huang, S.-Y. (2023). Improvement of cryo-EM maps by simultaneous local and non-local deep learning. Nature Communications 14, 3217. 10.1038/s41467-023-39031-1.

53. Jumper, J., Evans, R., Pritzel, A., Green, T., Figurnov, M., Ronneberger, O., Tunyasuvunakool, K., Bates, R., Žídek, A., and Potapenko, A. (2021). Highly accurate protein structure prediction with AlphaFold. nature 596, 583–589.

54. Emsley, P., and Cowtan, K. (2004). Coot: model-building tools for molecular graphics. Acta Crystallogr D Biol Crystallogr 60, 2126–2132. 10.1107/S0907444904019158.

55. Adams, P.D., Grosse-Kunstleve, R.W., Hung, L.W., Ioerger, T.R., McCoy, A.J., Moriarty, N.W., Read, R.J., Sacchettini, J.C., Sauter, N.K., and Terwilliger, T.C. (2002). PHENIX: building new software for automated crystallographic structure determination. Acta Crystallogr D Biol Crystallogr 58, 1948–1954. 10.1107/s0907444902016657.

56. Smart, O.S., Neduvelil, J.G., Wang, X., Wallace, B.A., and Sansom, M.S.P. (1996). HOLE: A program for the analysis of the pore dimensions of ion channel structural models. J Mol Graph Model 14, 354-&. Doi 10.1016/S0263-7855(97)00009-X.

57. Humphrey, W., Dalke, A., and Schulten, K. (1996). VMD: Visual molecular dynamics. J Mol Graph Model 14, 33–38. Doi 10.1016/0263-7855(96)00018-5.

58. Pettersen, E.F., Goddard, T.D., Huang, C.C., Couch, G.S., Greenblatt, D.M., Meng, E.C., and Ferrin, T.E. (2004). UCSF Chimera--a visualization system for exploratory research and analysis. J Comput Chem 25, 1605–1612. 10.1002/jcc.20084.

59. Pettersen, E.F. (2021). UCSF ChimeraX: structure visualization for researchers, educators, and developers. Protein Sci. 30. 10.1002/pro.3943.

60. Wu, E.L., Cheng, X., Jo, S., Rui, H., Song, K.C., Davila-Contreras, E.M., Qi, Y., Lee, J., Monje-Galvan, V., Venable, R.M., et al. (2014). CHARMM-GUI Membrane Builder toward realistic biological membrane simulations. J Comput Chem 35, 1997–2004. 10.1002/jcc.23702.

61. Lomize, M.A., Pogozheva, I.D., Joo, H., Mosberg, H.I., and Lomize, A.L. (2012). OPM database and PPM web server: resources for positioning of proteins in membranes. Nucleic Acids Res 40, D370–376. 10.1093/nar/gkr703.

62. Aho, N., Buslaev, P., Jansen, A., Bauer, P., Groenhof, G., and Hess, B. (2022). Scalable Constant pH Molecular Dynamics in GROMACS. J Chem Theory Comput 18, 6148–6160. 10.1021/acs.jctc.2c00516.

63. Buslaev, P., Aho, N., Jansen, A., Bauer, P., Hess, B., and Groenhof, G. (2022). Best Practices in Constant pH MD Simulations: Accuracy and Sampling. J Chem Theory Comput 18, 6134–6147. 10.1021/acs.jctc.2c00517.

64. Abraham, M.J., Murtola, T., Schulz, R., Páll, S., Smith, J.C., Hess, B., and Lindahl, E. (2015). GROMACS: High performance molecular simulations through multi-level parallelism from laptops to supercomputers. SoftwareX 1, 19–25.

65. Huang, J., Rauscher, S., Nawrocki, G., Ran, T., Feig, M., De Groot, B.L., Grubmüller, H., and MacKerell Jr, A.D. (2017). CHARMM36m: an improved force field for folded and intrinsically disordered proteins. Nature methods 14, 71–73.

66. Khan, H.M., MacKerell Jr, A.D., and Reuter, N. (2018). Cation-π interactions between methylated ammonium groups and tryptophan in the CHARMM36 additive force field. Journal of chemical theory and computation 15, 7–12.

67. Malde, A.K., Zuo, L., Breeze, M., Stroet, M., Poger, D., Nair, P.C., Oostenbrink, C., and Mark, A.E. (2011). An automated force field topology builder (ATB) and repository: version 1.0. Journal of chemical theory and computation 7, 4026–4037.

68. Hess, B., Bekker, H., Berendsen, H.J., and Fraaije, J.G. (1997). LINCS: a linear constraint solver for molecular simulations. Journal of computational chemistry 18, 1463–1472.

69. Darden, T., York, D., and Pedersen, L. (1993). Particle mesh Ewald: An N⋅ log (N) method for Ewald sums in large systems. The Journal of chemical physics 98, 10089–10092.

70. Berendsen, H.J., Postma, J.v., Van Gunsteren, W.F., DiNola, A., and Haak, J.R. (1984). Molecular dynamics with coupling to an external bath. The Journal of chemical physics 81, 3684–3690.

71. Bussi, G., Donadio, D., and Parrinello, M. (2007). Canonical sampling through velocity rescaling. The Journal of chemical physics 126.

72. Parrinello, M., and Rahman, A. (1981). Polymorphic transitions in single crystals: A new molecular dynamics method. Journal of Applied physics 52, 7182–7190.

73. Michaud-Agrawal, N., Denning, E.J., Woolf, T.B., and Beckstein, O. (2011). Software News and Updates MDAnalysis: A Toolkit for the Analysis of Molecular Dynamics Simulations. Journal of Computational Chemistry 32, 2319–2327. 10.1002/jcc.21787.

74. Adamson, L.S., Tasneem, N., Andreas, M.P., Close, W., Jenner, E.N., Szyszka, T.N., Young, R., Cheah, L.C., Norman, A., and MacDermott-Opeskin, H.I. (2022). Pore structure controls stability and molecular flux in engineered protein cages. Science Advances 8, eabl7346.

75. McKinney, W. (2010). Data structures for statistical computing in Python. In 1. pp. 51–56.

76. Harris, C.R., Millman, K.J., Van Der Walt, S.J., Gommers, R., Virtanen, P., Cournapeau, D., Wieser, E., Taylor, J., Berg, S., and Smith, N.J. (2020). Array programming with NumPy. Nature 585, 357–362.

77. Hunter, J.D. (2007). Matplotlib: A 2D graphics environment. Computing in science & engineering 9, 90–95.

78. Waskom, M.L. (2021). Seaborn: statistical data visualization. Journal of Open Source Software 6, 3021.

79. Virtanen, P., Gommers, R., Oliphant, T.E., Haberland, M., Reddy, T., Cournapeau, D., Burovski, E., Peterson, P., Weckesser, W., and Bright, J. (2020). SciPy 1.0: fundamental algorithms for scientific computing in Python. Nature methods 17, 261–272.

